# K_ir_2.1 modulation in macrophages sensitises dorsal root ganglion neurons through TNF secretion after nerve injury

**DOI:** 10.1101/2023.06.21.545843

**Authors:** Elena A. Konnova, Alexandru-Florian Deftu, Paul Chu Sin Chung, Guylène Kirschmann, Isabelle Decosterd, Marc R. Suter

## Abstract

Macrophages and satellite glial cells are found between injured and uninjured neurons in the lumbar dorsal root ganglia (DRG). We explored the mechanism of neuro-immune and neuron–glia crosstalk leading to hyperexcitability of DRG neurons. After spared nerve injury (SNI), CX3CR1^+^ resident macrophages became activated, proliferated and increased inward-rectifying potassium channel K_ir_2.1 currents. Conditioned medium (CM) by macrophages, obtained from DRG of SNI mice, sensitised small DRG neurons from naïve mice. However, treatment with CM from GFAP^+^ glial cells did not affect neuronal excitability. When subjected to this macrophage-derived CM, DRG neurons had increased spontaneous activity, current-evoked responses and voltage-gated Na_V_1.7 and Na_V_1.8 currents. Silencing K_ir_2.1 in macrophages after SNI prevented the induction of neuronal hyperexcitability from their CM. Blocking vesicular exocytosis or soluble tumour necrosis factor (TNF) in CM or interfering with the downstream intracellular p38 pathway in neurons, also prevented neuronal hyperexcitability. Blocking protein trafficking in neurons reduced the effect of CM, suggesting that the hyperexcitable state resulted from changes in Na_V_ channel trafficking. These results suggest that DRG macrophages, primed by peripheral nerve injury, contribute to neuron–glia crosstalk, Na_V_ channel dysregulation and neuronal hyperexcitability implicated in the development of neuropathic pain.

**Graphical abstract:** 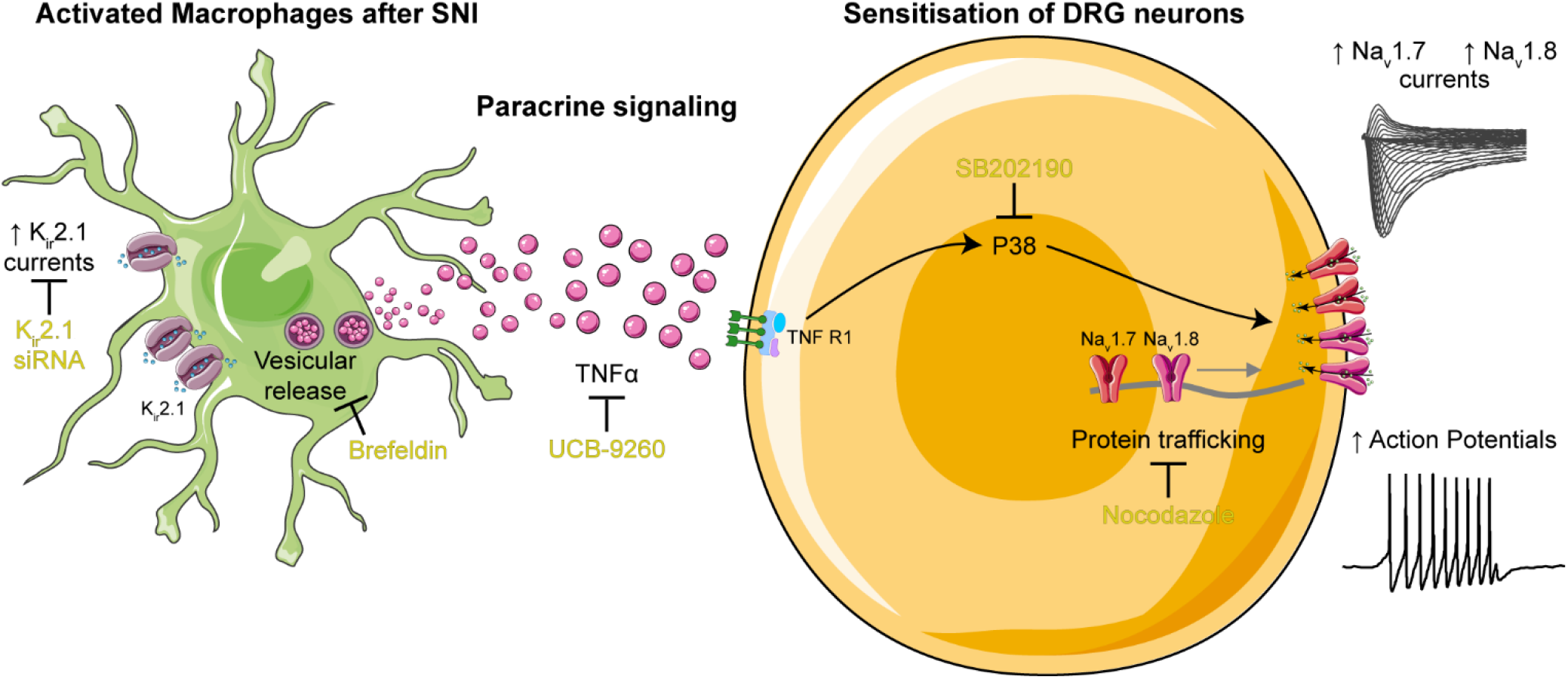

## Introduction

Nerve injury induces pathophysiological sensitisation of the pain pathway in the peripheral and central nervous system. Peripheral sensitisation increases action potential firing and decreases the firing threshold in primary sensory neurons, whose soma are located in dorsal root ganglia (DRG) (1, 2). In addition, ectopic activity is found at the neuroma site, along the axons and at the cell body level (3). Such hyperexcitability of a neuron can theoretically be achieved by dysregulation of a variety of ion channel at the expression and post-translational levels: for example, upregulation of voltage-gated sodium currents or downregulation of voltage-gated potassium currents can depolarise the resting membrane potential, reduce the action potential threshold and increase firing frequency (4). Nerve injury induces a downregulation of messenger RNA (mRNA) levels of most members of voltage-gated sodium channels (Na_V_) in DRG and the spinal nerve ligation (SNL) and spared nerve injury (SNI) mouse models (5, 6). Yet, Na_V_ currents, including Na_V_1.7 and Na_V_1.8 currents, are increased in ‘slow-conducting’ DRG neurons after SNI; these neurons are thought to be nociceptors. This increase in Na_V_ currents is mediated by the regulation of internalisation of the channels from the cell surface membrane (7).

In the SNI model, injured and uninjured neurons are in close proximity within the same DRG, but the ligated and spared axons are segregated (6). Alongside primary sensory neurons, DRG also host macrophages (8–10) and satellite glial cells (SGC) that play protective, supporting and modulatory roles (11–13). This anatomical configuration facilitates cell–cell interactions between neurons and surrounding immune and glial cells, and mediates the cross-sensitisation of uninjured neurons after SNI (2, 9, 12, 14). Nerve injury induces immune and glial reactions that contribute to the establishment and maintenance of neuropathic pain if it remains unresolved (8, 12, 14–16). Neuroinflammation sensitises neurons through the secretion of pro-inflammatory cytokines from immune cells, such as macrophages, that can directly increase the excitability of neurons by acting on receptors, activating intracellular signalling and modulating neuronal ion channels (14, 17, 18)

In the periphery, macrophages activate, proliferate and change state to release pro-inflammatory cytokines in response to nerve injury at the site of injury, along the axons in the nerve, and at the level of the soma of primary sensory neurons in the DRG (8, 14, 16). Preventing activation or depleting peripheral macrophages rescues the hypersensitivity after nerve injury (9, 19, 20). Intravenous administration of minocycline or depleting agents can act on nerve and DRG macrophages (20). Depending on the experimental approaches, some studies have shown that targeting infiltrating macrophages in the nerve can attenuate nerve-injury-induced hypersensitivity (21). Other studies have highlighted the significant contribution of resident macrophages from DRG to the initiation and maintenance of hypersensitivity and have demonstrated that depletion of nerve macrophages is insufficient to eliminate hyperalgesia (20).

We designed this *ex vivo* study to investigate the process of sensitisation of uninjured neurons via neuro-immune and neuron–glia crosstalk occurring specifically within the DRG. We studied the effect of the paracrine signalling of DRG CX3CR1^+^ macrophages and glial fibrillary acid protein (GFAP)^+^ SGC from mice after SNI, by transferring the culture medium from these cells to DRG neurons from naïve mice. We aimed to characterise DRG macrophages after SNI and to elucidate the mechanism behind the sensitisation of neurons by the macrophage medium. Finally, we measured the impact of the sensitisation on the excitability and currents from Na_V_ channels in the treated small DRG neurons, which are regarded as nociceptors.

## Results

### Culture medium conditioned by fluorescence-activated cell sorting (FACS) of CX3CR1^+^ cells from DRG 7 days after SNI sensitise primary sensory neurons

To investigate the neuro-immune and neuron–glia crosstalk in DRG, we used the CX3CR1-GFP transgenic mouse line, which labels CX3CR1^+^ cells, considered to be macrophages, and the hGFAP-CFP transgenic mouse line, which labels the GFAP^+^ glial cells in DRG (11, 22, 23). To explore the role of paracrine signalling in the sensitisation of DRG neurons after peripheral nerve injury, we sorted CX3CR1^+^ and GFAP^+^ cells by FACS from the ipsilateral and contralateral L3–L5 DRG of transgenic mice 7 days after SNI. We cultured them *ex vivo* for 3 days to allow signalling molecules to be secreted into the culture medium. The collected culture medium is considered to be conditioned medium (CM) by CX3CR1^+^ or GFAP^+^ cells after SNI. We then applied CM to the culture of dissociated DRG neurons from naïve mice to study the effect of CM on neuronal excitability (Figure 1A).

**Figure 1:**
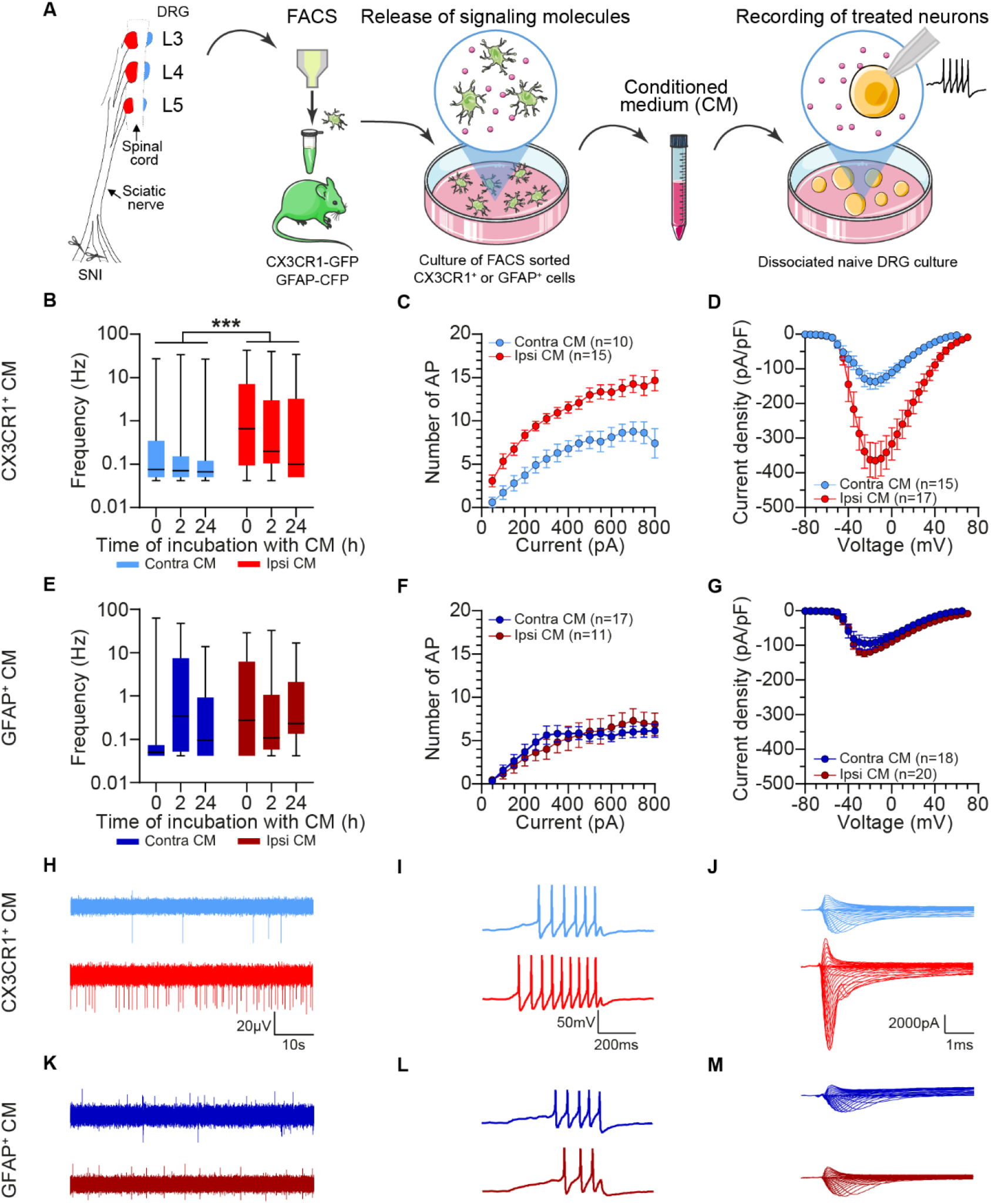
Effect of CX3CR1^+^ or GFAP^+^ cell conditioned medium (CM) from mice 7 days after spared nerve injury (SNI) on dorsal root ganglion (DRG) neurons. (A) Schematic representation of the experimental design: ipsilateral and contralateral L3–L5 DRG from CX3CR1-GFP or GFAP-CFP transgenic mice were collected 7 days after SNI, dissociated and submitted to fluorescence-activated cell sorting. The sorted cells were then cultured for 3 days and the collected culture medium is considered CM. CM was then applied on DRG neurons from naïve mice and the current was recorded by multielectrode array (MEA) with patch clamping. CM from DRG ipsilateral to the nerve injury is the ipsilateral CM and the control is CM from DRG contralateral to the nerve injury. (B, E) MEA recordings of spike frequency of DRG neuron cultures treated for <10 min, 2 hours, and 24 hours with CM from CX3CR1^+^ or GFAP^+^ cells (two-way analysis of variance [ANOVA], treatment type ***p <0.001 [B], p >0.05 [E]). Example of MEA raw traces after 2-hour treatment with CM from CX3CR1^+^ (H) or GFAP^+^ cells (K). (C, F) Current-evoked response of small DRG neurons treated for 2 hours with CM from ipsilateral or contralateral CX3CR1^+^ or GFAP^+^ cells (two-way ANOVA with repeated measures, treatment p <0.001 [C], p >0.05 [F]). (I, L) Representative traces at 250-pA steps of DRG neurons treated for 2 hours with CM from CX3CR1^+^ (I) or GFAP^+^ (L) cells. (D, G) Total voltage-gated sodium channel (Na_V_) current density of DRG neurons treated for 2 hours with CM from ipsilateral or contralateral CX3CR1^+^ or GFAP^+^ cells (two-way ANOVA with repeated measures, treatment p <0.001 [D], p >0.05 [G]). (J, M) Example voltage clamp traces from DRG neurons treated for 2 hours with CM from CX3CR1^+^ (J) or GFAP^+^ (M) cells.

We recorded field potential of the dissociated DRG neurons after CM treatment using multielectrode array (MEA) to assess spontaneous neuronal activity. CM from ipsilateral CX3CR1^+^ cells after SNI increased the spiking frequency of DRG neurons compared with CM from contralateral CX3CR1^+^ cells. The effect was present immediately and at 2 and 24 hours after incubation with CM (Figure 1B, H). Whole-cell current patch-clamp recordings demonstrated an increase in the action potential frequency of small DRG neurons treated with CM from ipsilateral CX3CR1^+^ cells when we delivered increasing ramp currents compared with CM from contralateral CX3CR1^+^ cells (Figure 1C, I). Voltage clamp recordings revealed that DRG neurons treated with CM from ipsilateral CX3CR1^+^ cells had significantly larger amplitude of total Na_v_ current density compared with DRG neurons treated with CM from contralateral CX3CR1^+^ cells (Figure 1D, J). We observed none of these differences when using CM from GFAP^+^ cells (Figure 1E–G, K–M).

### Changes in CX3CR1^+^ cells can contribute to DRG neuron hyperexcitability after SNI

Transient proliferation of CX3CR1^+^ cells, identified by co-localisation with the Ki67 marker, peaked at 2 and 4 days after SNI in L3 and L4 DRG (Figure 2A, C). In the same DRG at days 4 and 7 after SNI, the number of CX3CR1^+^ cells increased compared with naïve and sham-operated mice. We did not observe these changes in L5 DRG. We also found an increase in Iba1 protein, another marker of DRG macrophages, 7 days after SNI compared with sham surgery (Figure 2D).

**Figure 2:**
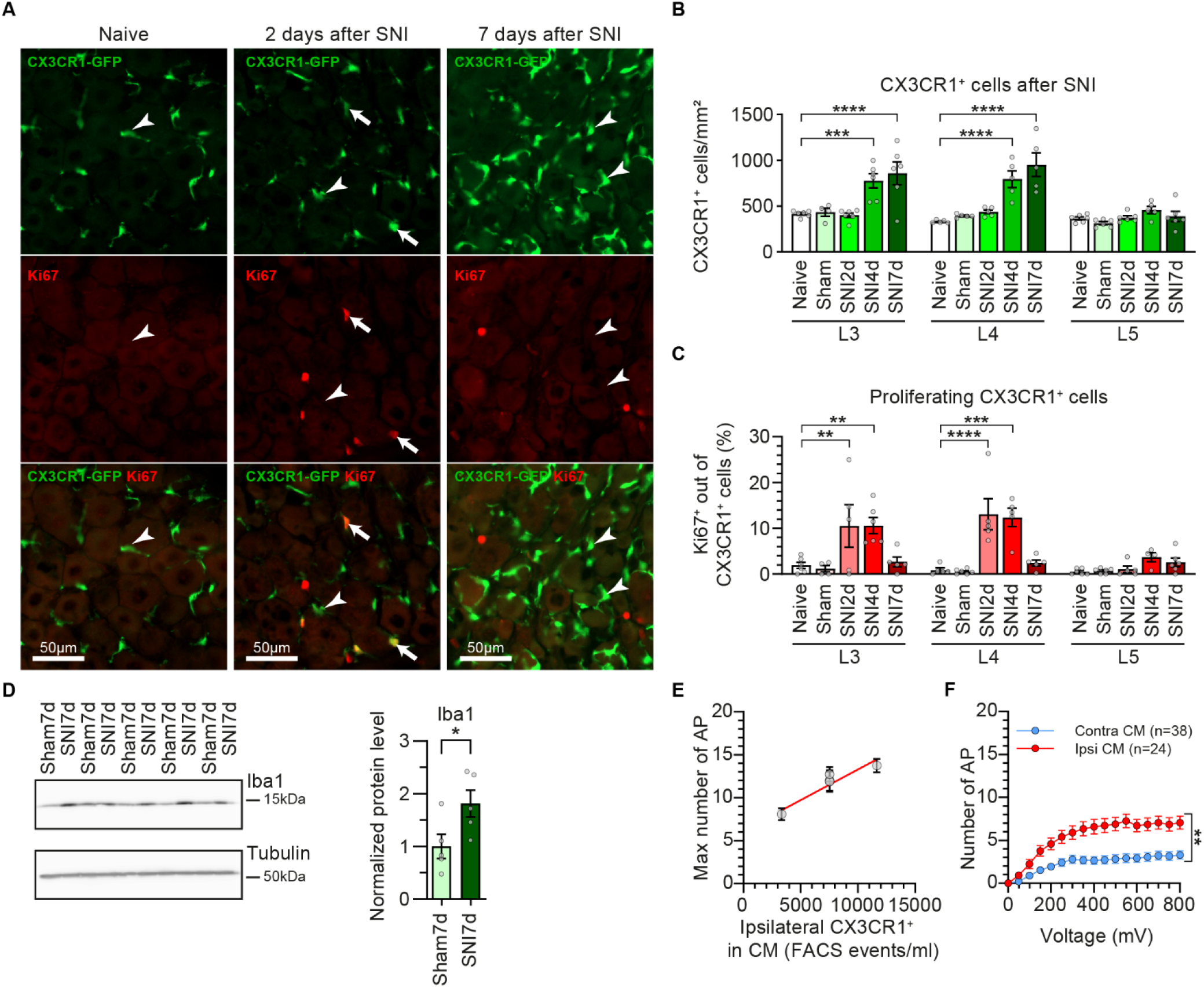
Proliferation and state of ipsilateral CX3CR1^+^ cells after spared nerve injury (SNI) affect the potency of conditioned medium (CM). (A) Representative images of L4 dorsal root ganglion (DRG) sections from naïve mice, or mice 2 or 7 days after SNI. Native GFP signal form CX3CR1-GFP mice (green) and immunohistochemistry for the marker of proliferation Ki67 (red). The arrows indicate CX3CR1^+^ cells with co-localisation of Ki67; the arrowheads indicate CX3CR1^+^ cells that are not co-localised. (B) Quantification of the number of CX3CR1^+^ cells in ipsilateral L3, L4 and L5 DRG sections from naïve mice; sham-operated mice (7 days); or mice 2, 4 and 7 days after SNI. (C) Percentage of proliferating CX3CR1^+^ cells, identified by the co-localisation with Ki67 (two-way analysis of variance [ANOVA], interaction p <0.0001 [B], p <0.001 [C], with *post hoc* Dunnett’s comparisons to naïve mice, n = 6). (D) Western blot for Iba1 of ipsilateral L3–L5 DRG from mice at 7 days after SNI or Sham surgery (n = 5). (E) Effect of the number of fluorescence-activated cell sorting events on the excitability of DRG neurons treated with the CM for each individual experiment (n = 5). The maximum number of action potentials in current clamp correlates proportionally to the concentration of ipsilateral CX3CR1^+^ cells used for the CM (Pearson’s correlation r = 0.935, R² = 0.874, p <0.05). (F) Current-evoked response of small DRG neurons treated for 2 hours with CM from ipsilateral or contralateral CX3CR1^+^ cells, which was prepared with an equal concentration of sorted cells (two-way ANOVA, treatment p <0.0001). n = number of recorded neurons.

The number of CX3CR1^+^ events sorted by FACS from ipsilateral DRG was higher than from the contralateral side at 7 days after SNI (Supplemental Table 1). To understand whether the sensitisation effect was impacted by the number of CX3CR1^+^ events, we performed a correlation analysis across five independent experiments, in which we treated DRG neurons with CM from different numbers of CX3CR1^+^ events sorted by FACS. We found a positive correlation between the number of ipsilateral CX3CR1^+^ events used to prepare the CM and the maximum number of action potentials evoked in current clamp (Figure 2E).

We controlled for the number of CX3CR1^+^ cells by plating an equal density of ipsilateral and contralateral CX3CR1^+^ cells during CM preparation, to determine whether a phenotypic change of CX3CR1^+^ cells underlies the sensitisation effect (Supplemental Table 1). After normalising the cell number, CM from ipsilateral CX3CR1^+^ cells still induced a greater action potential response compared with CM from contralateral CX3CR1^+^ cells (Figure 2F). These results suggest that the phenotype of CX3CR1^+^ cells impacts the excitability state of DRG neurons.

### CX3CR1**^+^** cells alter their state and downregulate major histocompatibility complex class II (MHC II)

We aimed to characterise the phenotype of DRG CX3CR1^+^ cells after SNI. We performed liquid chromatography with tandem mass spectrometry (LC-MS/MS) on bulk samples of either ipsilateral or contralateral CX3CR1^+^ cells sorted by FACS at 7 days after SNI to identify changes in the proteome (Figure 3D). Ipsilateral CX3CR1^+^ cells presented upregulation of cyclic ADP ribose hydrolase (CD38) and tissue transglutaminase (Tgm2), and downregulation of arginase 1 (Arg1) and HLA class II histocompatibility antigen gamma chain (CD74), among other significantly regulated proteins (Supplemental Table 2). CD74 invariant chain protein mediates the assembly of MHC II, which is a commonly used marker of lipopolysaccharide (LPS)-induced M1-activation state (24). We observed two populations of CX3CR1^+^ cells after SNI (presented as mean ± standard error of the mean [SEM]): 62% ± 3.6% were MHC II^+^ and 38% ± 3.6% were MHC II^-^ in naïve mice. Seven days after SNI, the proportion of MHC II^-^ CX3CR1^+^ cells increased to 56.4% ± 4.5% in the L3 and L4 DRG. The change in proportion occurred due to the increase in MHC II^-^ CX3CR1^+^ cells, while MHC II^+^ CX3CR1^+^ cells remained stable in number after SNI (Figure 3A–C).

**Figure 3:**
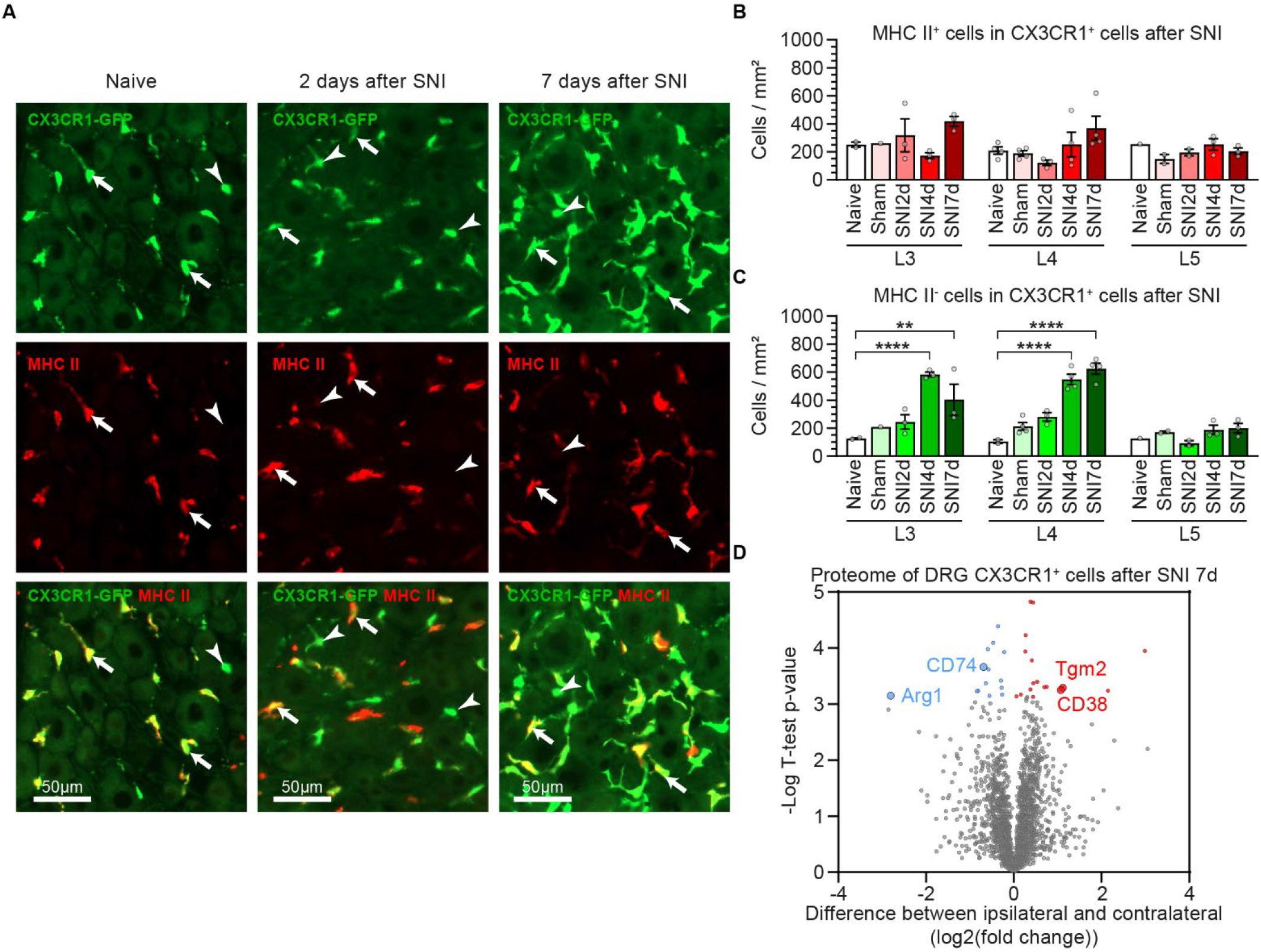
Activation profile of dorsal root ganglion (DRG) CX3CR1+ cells after spared nerve injury (SNI). (A) Representative images of L4 DRG sections from naïve mice or CX3CR1-GFP transgenic mice 2 or 7 days after SNI. Native GFP (green) and immunohistochemistry for MHC II (red). The arrows indicate CX3CR1+ cells with co-localisation of MHC II; the arrowheads indicate CX3CR1+ cells that are not co-localised. (B) The number of MHC II+ CX3CR1+ cells and (C) the number of MHC II-CX3CR1+ cells in the L3, L4 and L5 DRG of naïve mice; sham-operated mice; and mice 2, 4 and 7 days after SNI (two-way analysis of variance [ANOVA], interaction p >0.05 [B], p <0.01 with post hoc Dunnett’s compared with naïve [C], n = 6). (D) Proteome differences between CX3CR1+ cells (from fluorescence-activated cell sorting) from ipsilateral and contralateral L3–L5 DRG of mice 7 days after SNI, analysed by liquid chromatography coupled to tandem mass spectrometry. Significantly upregulated proteins are shown in red, and significantly downregulated proteins are shown in blue (n = 3).

### CX3CR1**^+^** cells upregulate the K_ir_2.1 current, which is necessary for them to sensitise DRG neurons

The electrophysiological state of non-excitable immune cells such as microglia has previously been shown to regulate their function after SNI (25). Therefore, we characterised the currents of CX3CR1^+^ cells from an *ex vivo* culture of dissociated DRG after SNI. Ipsilateral CX3CR1^+^ cells were larger, as indicated by their larger capacitance compared with contralateral CX3CR1^+^ cells (Figure 4A). The resting membrane potential measured in current clamp mode was not significantly different between the two groups (ipsilateral macrophages −56.4 ± 3.4 mV, contralateral macrophages −57.2 ± 4.1 mV, Figure 4B). CX3CR1^+^ cells had small inward currents, but no outward currents. Ipsilateral DRG CX3CR1^+^ cells had an increased inward potassium current density 7 days after SNI compared with contralateral DRG CX3CR1^+^ cells (Figure 4C, D). Those inward potassium currents were inhibited by the K_ir_2.1 inhibitor ML133, indicating a significant increase in K_ir_2.1 currents in ipsilateral CX3CR1^+^ cells after SNI. We found no significant difference in the rates of proliferation and the increase in K_ir_2.1 currents of macrophages from male and female mice (data not shown).

**Figure 4:**
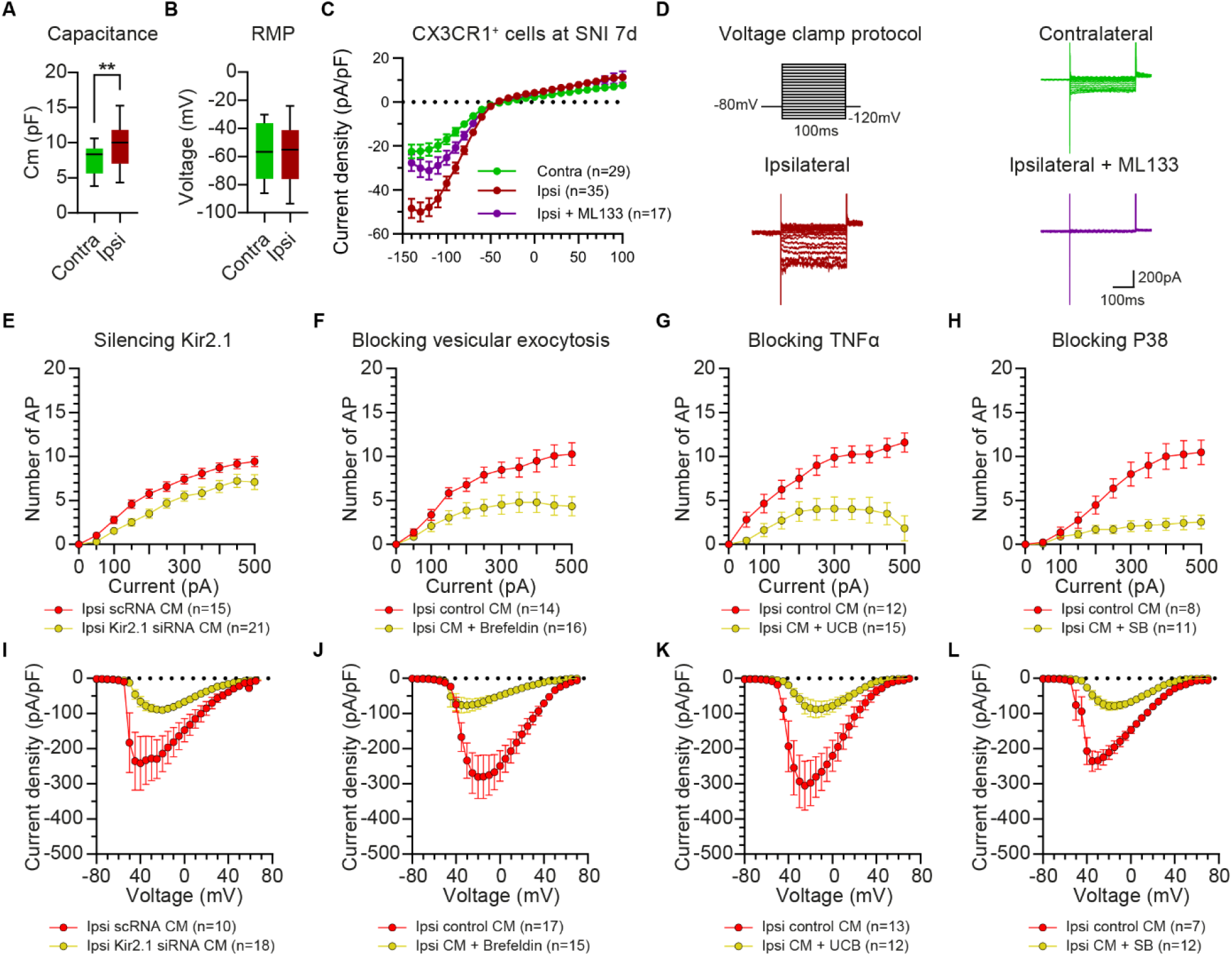
Increase in Kir2.1 currents in CX3CR1+ dorsal root ganglion (DRG) cells after spared nerve injury (SNI) are necessary for paracrine sensitisation of DRG neurons via the tumour necrosis factor (TNF)–p38 pathway. (A) Capacitance of patched CX3CR1+ cells. (B) Resting membrane potential (RMP) of CX3CR1+ cells in standard extracellular solution (contra n = 21, ipsi n = 31). (C) Current densities of CX3CR1+ cells 7 days after SNI in response to the step voltage clamp protocol, recorded in extracellular solution with 20 mM KCl. ML133 (50 μM) added in the extracellular solution was used to block Kir2.1 currents (two-way analysis of variance [ANOVA], interaction p <0.0001, with post hoc Tukey’s comparisons: ipsi vs contra and ipsi vs ipsi + ML133 p <0.0001, contra vs ipsi + ML133 p >0.05). (D) Example of voltage clamp traces from DRG CX3CR1+ cells after SNI. (E) Current-evoked response of small DRG neurons treated with conditioned medium (CM) from ipsilateral macrophages after SNI incubated with scramble RNA (scRNA) or silencing RNA (siRNA) for Kir2.1. Current-evoked response of small DRG neurons treated with CM from ipsilateral CX3CR1+ cells, with or without pharmacological blockers: brefeldin to block vesicular release in macrophages (F), UCB-9260 to block TNF signalling from macrophages to neurons (G) and SB202190 to block intracellular p38 signalling in DRG neurons (H) (two-way ANOVA, treatment p <0.0001 [E], p <0.01 [F], p <0.001 [G, H]; interactions p <0.0001 [E–H]). (I) Total NaV current density of small DRG neurons treated with CM from ipsilateral CX3CR1+ cells after SNI incubated with scRNA or Kir2.1 siRNA. Total NaV current density of small DRG neurons treated with CM from ipsilateral CX3CR1+ cells, with or without pharmacological blockers: 50 μM brefeldin (J), 10 μM UCB-9260 (K) and 10 μM SB202190 (L) (two-way ANOVA, treatment p <0.01 [I, K], p <0.05 [J], p <0.0001 [L]; interactions p <0.0001 [I–L]) n = number of recorded neurons.

Silencing K_ir_2.1 in CX3CR1^+^ cells during the preparation of CM by applying K_ir_2.1 silencing RNA (siRNA) for 24 hours prevented the sensitisation effect compared with using scramble RNA (scRNA): the frequency of current-evoked action potentials and the total Na_V_ current densities was decreased compared with the scRNA-treated control group (Figure 4E, I).

### CX3CR1**^+^** cells sensitise DRG neurons through the tumour necrosis factor (TNF)–p38 pathway

Our experimental design prevents cell–cell communication through direct contact that would occur in a co-culture of macrophages and neurons. Instead, it involves the transfer of CM that would contain soluble signalling molecules released by CX3CR1^+^ cells. Blocking vesicular release of CX3CR1^+^ cells for 24 hours prior to the preparation of CM using 50 μM brefeldin on ipsilateral CX3CR1^+^ cells from SNI mice prevented the sensitisation effect of CM. DRG neurons had a reduced current-evoked response and lower total Na_V_ current density (Figure 4F, J). These results suggest that CX3CR1^+^ cells are capable of sensitising DRG neurons via paracrine signalling.

We investigated the involvement of soluble TNF in the sensitisation of DRG neurons by CM from CX3CR1^+^ cells. DRG cultures from naïve mice were treated with CM from CX3CR1^+^ cells containing 10 μM UCB-9260, a synthetic blocker that directly binds to TNF to prevent its binding to the TNF receptor. Blocking TNF signalling in CM from ipsilateral CX3CR1^+^ cells prevented DRG neuron hyperexcitability, suggesting that TNF participates in the sensitisation effect of CX3CR1^+^ cells. The number of action potentials evoked by current stimulation and the total Na_V_ current density were significantly reduced by the addition of the TNF blocker to CM from ipsilateral CX3CR1^+^ cells (Figure 4G, K).

Previous studies have shown that modulation of Na_V_ currents in DRG neurons by extracellular TNF is mediated by the downstream p38 response (26, 27). To determine whether the sensitisation effect of CM is mediated by the TNF–p38 pathway, we added 10 μM SB202190, a p38 blocker, to CM from ipsilateral CX3CR1^+^ cells. Blocking p38 also resulted in fewer current-evoked action potentials during current clamp recordings, and a decreased total Na_V_ current density in voltage clamp recordings, compared with CM from ipsilateral CX3CR1^+^ cells without the blocker (Figure 4H, I). Thus, we demonstrated that TNF and p38 activation are necessary to mediate the sensitisation of DRG neurons.

### CM from CX3CR1**^+^** cells induces an increase in the Na_V_1.8 and Na_V_1.7 current densities

To distinguish between Na_V_ channel subtypes, we electrically blocked Na_V_1.9, then perfused extracellular solution containing protoxin II (ProTxII), a Na_V_1.7 blocker, followed by tetrodotoxin (TTX) to block all remaining Na_V_ channels, except TTX-resistant Na_V_1.8 (Figure 5M). The patched small DRG neurons had the largest contribution from Na_V_1.7 and Na_V_1.8, while other TTX-sensitive currents densities were very low at their peak amplitude (57.7 ± 7.4 pA/pF). The amplitude of Na_V_1.7 and Na_V_1.8 current densities were increased in DRG neurons sensitised by CM from ipsilateral CX3CR1^+^ cells compared with CM from contralateral CX3CR1^+^ cells (Figure 5E, I).

**Figure 5:**
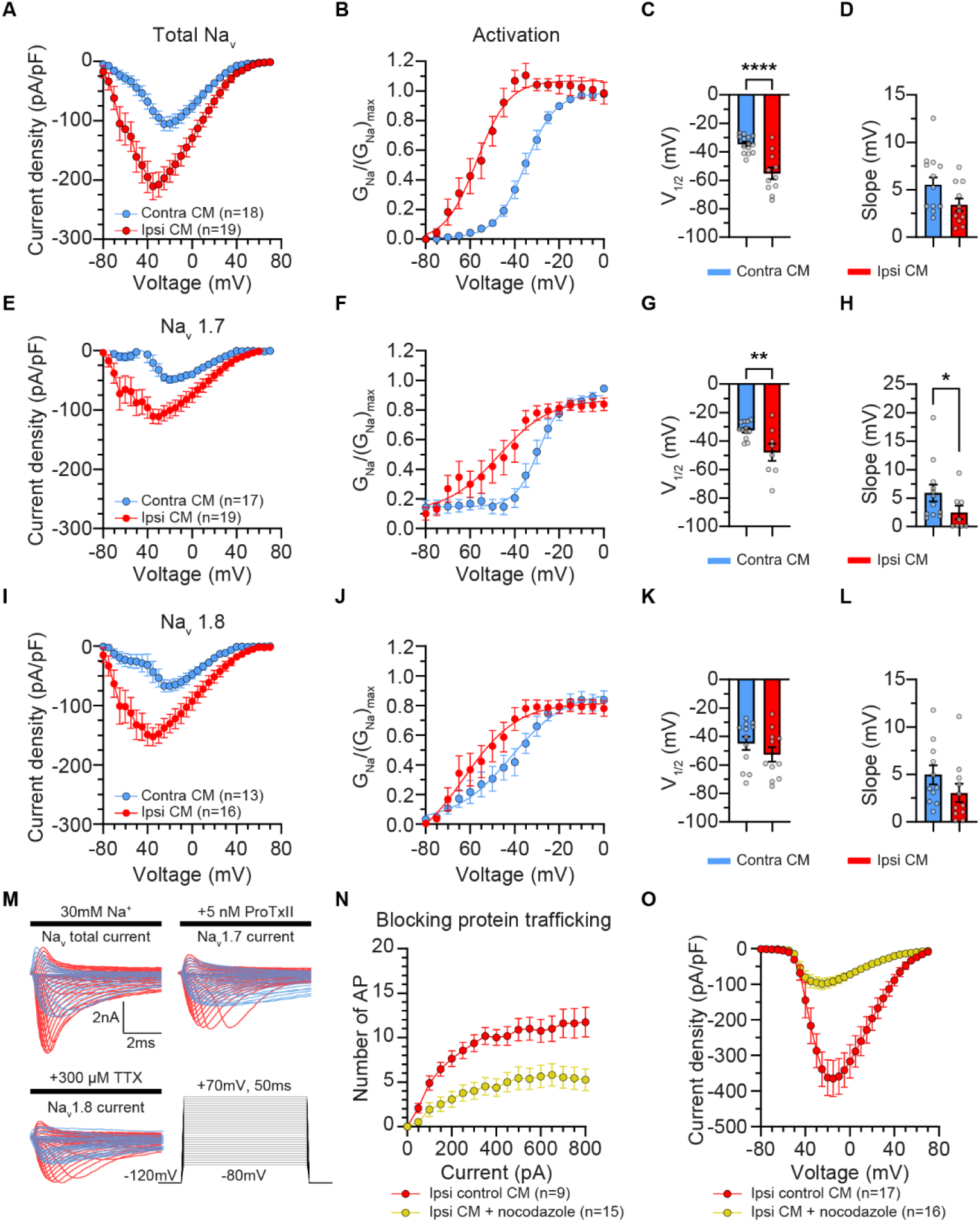
Effect of conditioned medium (CM) from CX3CR1+ cells on voltage-gated sodium currents of dorsal root ganglion (DRG) neurons. Total NaV current densities in DRG neurons treated with CM from ipsilateral or contralateral CX3CR1+ cells 7 days after spared nerve injury (SNI). (A–D) Total NaV current was recorded in absence of blocker and an electrical protocol was used to inactivate the NaV1.9 current. (E–H) The NaV1.7 current was isolated by subtracting the total NaV current following application of the NaV1.7 blocker Protoxin II (ProTxII). (I–L) The NaV1.8 current was obtained by blocking the remaining tetrodotoxin-sensitive (TTX-S) current by applying TTX (two-way analysis of variance [ANOVA], treatment p <0.0001 [A, E, I]; interaction p <0.0001 [A, E, I]). Steady-state activation curves of total NaV (B), NaV1.7 (F) and NaV1.8 (J) currents in small DRG neurons treated with CM from ipsilateral or contralateral CX3CR1+ cells. The half-maximal activation (V½) (C, G, K) and slope (D, H, L) were calculated for each activation curve. (M) Example traces of total NaV current, after ProTxII and after TTX application in small DRG neurons previously treated with CM from ipsilateral (red) or contralateral (blue) CX3CR1+ cells. (N) Current-evoked response of small DRG neurons treated with CM from ipsilateral CX3CR1+ cells, with or without 10 μM nocodazole to block trafficking of proteins to the membrane surface in neurons (two-way ANOVA, treatment p <0.001; interactions p <0.0001). (O) Total NaV current density of small DRG neurons treated with CM from ipsilateral macrophages, with or without 10 μM nocodazole (two-way ANOVA, treatment p <0.001; interactions p <0.0001). n = number of recorded neurons.

The observed increase in Na_V_ current densities could result from several mechanisms leading to an accumulation of Na_V_ channels at the surface membrane or due to changes in channel biophysical properties. Indeed, the steady state activation of Na_V_1.7 was shifted by CM from ipsilateral CX3CR1^+^ cells, indicated by the lower V_½_ (Figure 5G). The inactivation properties of total Na_V_, Na_V_1.7 and Na_V_1.8 remained unchanged by CM from ipsilateral CX3CR1^+^ cells (Supplemental Figure 1). We then added 10 μM nocodazole to CM during treatment of DRG culture from naïve mice to block transport of proteins along the microtubules and to prevent the trafficking of Na_V_ channels to the membrane. Nocodazole prevented the increase in action potentials and in total Na_V_ current densities induced by CM from ipsilateral CX3CR1^+^ cells, indicating that protein trafficking is necessary for the sensitisation effect (Figure 5N, O).

## Discussion

In this study, we have described a new neuro-immune interaction in the DRG in the SNI model of neuropathic pain. After peripheral nerve injury, CX3CR1^+^ DRG cells proliferate, modify their phenotype and induce nociceptor hyperexcitability. DRG neurons sensitised by CM from SNI-primed macrophages show increased spontaneous activity, increased current-evoked action potentials and greater Na_v_1.7 and Na_v_1.8 current density. Blocking the K_ir_2.1 current or vesicular exocytosis in CX3CR1^+^ cells or blocking the TNF–p38 signalling pathway or protein trafficking in the nociceptors reduces the hyperexcitability.

Peripheral nerve injury induces the proliferation of resident DRG macrophages (9, 10, 28). We found that CX3CR1^+^ macrophages undergo a transient phase of proliferation, which subsided 7 days after SNI. By that time, the CX3CR1^+^ macrophage population had doubled. We found that the number of CX3CR1^+^ cells and their state determined the capacity of their CM to sensitise DRG neurons. The activation of macrophages was sufficient to make them release sensitising factors, while their proliferation amplified the effect.

CX3CR1^+^ macrophages in the DRG modified their phenotype after SNI. At 7 days after SNI, bulk analysis of the DRG macrophage proteome showed conflicting regulation of M1 or M2 markers. The M1/M2 nomenclature describes the state of *in vitro*–activated macrophages by, respectively, LPS/interferon gamma (IFN-γ) or interleukin 4 (IL-4)/IL-10 (29, 30). Multiple states of CX3CR1^+^ macrophages may be hidden within our bulk analysis. Nonetheless, other studies have demonstrated that the phenotype of an individual DRG macrophage after nerve injury does not fit into the M1/M2 paradigm (31). We observed MHC II^-^ CX3CR1^+^ macrophages, which increase in number, and MHC II^+^ CX3CR1^+^ macrophages, which remain stable after SNI. Other studies have found an increase in overall MHC II expression in DRG after spinal nerve transection, sciatic nerve transection and chronic constriction injury of the sciatic nerve in rats (32, 33). This discrepancy could be explained by the expression of MHC II in other immune cell populations and the fact that macrophage regulation is tissue, species, disease and time-course specific (30, 31).

In the SNI model, ligated and spared afferents are not mixed in the sciatic nerve, but their somas reside in close proximity to each other within the same DRG. Therefore, the DRG is an important location where macrophages primed by injured neurons can sensitise neighbouring uninjured neurons. The proliferation and activation of CX3CR1^+^ macrophages take place in L3 and L4 DRG, where most injured neurons are present following SNI (6), but not in L5 DRG, which mainly contain uninjured neurons from the sural nerve. Evidence from other studies show that signalling from injured neurons, such as CSF1, triggers the activation of DRG macrophages after SNI (9, 34). In this study, we showed that macrophages primed by SNI release pro-inflammatory factors that sensitise DRG neurons. Through this mechanism, the sensitisation of uninjured neurons may be the origin of the hypersensitivity of the spared sural nerve after SNI (2, 6).

Beyond commonly used markers, we further characterised the electrophysiological changes in the macrophage phenotype. We demonstrated that CX3CR1^+^ macrophages from ipsilateral DRG after SNI had increased K_ir_2.1 current density. K_ir_2.1 is an inward-rectifying potassium channel that is expressed in a wide variety of excitable and non-excitable cells, including macrophages and microglia (25, 35, 36). Manipulating the electrophysiological state of macrophages may be a tool to modulate their state and function (37). A recent study using a macrophage cell line demonstrated that *in vitro* knockdown or blockade of K_ir_2.1 polarises the cells towards an anti-inflammatory profile (36). However, in the context of cancer research, tumour-associated macrophages with an M2 profile are polarised towards an M1-like state after K_ir_2.1 inhibition (38). We have previously shown the upregulation of K_ir_2.1 currents in spinal microglia and their necessity for the proliferative and pro-nociceptive microglial profile in the SNI model (25). *In vivo* intrathecal injection of the K_ir_2.1 inhibitor ML133 reduced the proliferation of spinal microglia and attenuated hypersensitivity after SNI. Here, we demonstrated that by silencing K_ir_2.1 in CX3CR1^+^ macrophages from SNI mice, we can reduce the sensitisation of DRG neurons.

Some neuro-immune interactions in the context of pain are sex dependent. Activation of spinal cord microglia is necessary for the development of chronic pain hypersensitivity in male mice, but it is not required for female mice, which rely on the activity of T lymphocytes (39). However, development of mechanical hypersensitivity to TNF occurs in male and female mice (40). In this study, we used male and female mice and saw no differences between the sexes in the proliferation response, MHC II expression and upregulation of K_ir_2.1 currents in DRG macrophages. Using both sexes to prepare macrophage CM, we found a consistent effect on neuronal excitability. As we have not found significant evidence for sex differences in the DRG macrophage characterisation and the *ex vivo* sensitisation experiments, we did not pursue further investigations regarding the impact of sex.

In our experimental setting, we sensitised DRG neurons using CM, implying a soluble pro-nociceptive factor released by activated macrophages. Therefore, we studied the involvement of TNF, a pro-nociceptive cytokine, because it is well known to be upregulated by macrophages after nerve injury (41–43). Its administration to the sciatic nerve or DRG is sufficient to induce neuropathic pain (44, 45) and direct application of TNF to DRG neurons from naïve mice elicits neuronal discharges (45). Endogenously, soluble TNF is released from vesicles after cleavage of its transmembrane form and can bind to TNFR1 and TNFR2 receptors (46). We blocked TNF in macrophage CM by using a synthetic drug that directly binds to the molecule, rather than the receptor. This blockade prevented the sensitisation of DRG neurons. Therefore, we confirmed that soluble TNF released from DRG macrophage CM is a major sensitising factor in the SNI model. However, the contribution of other cytokines and signalling molecules is not excluded.

Previous studies have described the pathway and impact of TNF on Na_V_ channels. Application of TNF to DRG neurons upregulates TTX-resistant Na_V_ currents through TNFR1 (26). TNFR1 activation leads to the intracellular phosphorylation of p38, which, in turn, modulates Na_V_ currents (26). Researchers have shown that the TNF–p38 pathway is necessary to induce hypersensitivity in a nerve injury model (27). By blocking p38 in DRG neurons, we similarly prevented sensitisation of DRG neurons by CM from CX3CR1^+^ macrophages. These results suggest that sensitisation of DRG neurons by macrophages after SNI depends on soluble TNF and its downstream p38 activation.

We studied the Na_V_ currents downstream of the TNF–p38 pathway. Na_V_1.7 and Na_V_1.8 are dominant Na_V_ currents in the small DRG neurons considered to be nociceptors (47). We showed that treatment with CM from CX3CR1^+^ macrophages from SNI mice was sufficient to increase Na_V_1.7 and Na_V_1.8 currents in DRG neurons from naïve mice. Na_V_1.7 and Na_V_1.8 currents are increased in the small DRG neurons of SNI mice, but it remains unclear whether this effect occurs in injured or uninjured neurons (47). Injured DRG show the greatest sensitivity to direct application of TNF, but uninjured in adjacent DRG in the SNL model also an increased compared to naïve DRG (45). The sensitivity to TNF may be modulated by the changes in TNFR1 and TNFR2 expression after nerve injury (48). The increase in Na_V_1.7 and Na_V_1.8 currents in DRG neurons following CM treatment was accompanied by changes in gating properties of Na_V_1.7 but not Na_V_1.8 channels. We observed a leftward shift in the activation curve of the Na_V_1.7 channels to lower voltage, with the V_½_ approaching the resting membrane potential of DRG neurons. Consistently, other studies have shown that direct application of TNF on DRG neurons does not change the gating properties of TTX-resistant Na_V_ channels, but it does induce a shift in TTX-sensitive Na_V_ currents (26, 49).

Finally, we demonstrated that trafficking of Na_V_ channels to the surface membrane is necessary to mediate the sensitisation of the treated DRG neurons. Treatment of DRG neurons with nocodazole prevents the transport of Na_V_ channels, among other proteins, to the surface membrane by destabilising microtubules (50). Nocodazole prevented CM-mediated sensitisation of DRG neurons by and maintained low Na_V_ currents. Regulation of trafficking of Na_V_ channels at the surface membrane is a mechanism that reconciles the apparent downregulation of mRNA levels of most Na_V_ members, yet the upregulation of Na_V_ currents in DRG neurons after SNI (6, 47, 51). In addition, other mechanisms, such as the regulation of internalisation of Na_V_ channels from the surface membrane, can modulate Na_V_ currents. Previous research found the reduction of internalisation of Na_V_1.7 from the surface membrane after SNI via NEDD4.2, a ubiquitin protein ligase (47). TNF or macrophage CM treatment may impact Na_V_ channel trafficking to and from the surface membrane.

We conclude that resident DRG CX3CR1^+^ macrophages after SNI proliferate and modify their properties, which enables them to sensitise DRG neurons from naïve mice. Resident DRG macrophages likely become activated following signals from injured neurons in L3 and L4 DRG. In turn, activated macrophages sensitise neighbouring uninjured neurons. Thus, the hypersensitivity of the uninjured sural nerve may be mediated by the mechanism of sensitisation in the DRG. The sensitisation of small nociceptive DRG neurons after SNI is largely mediated by the TNF–p38 pathway that modulates the amplitude of Na_V_1.7 and Na_V_1.8 currents and gating properties of Na_V_1.7. The upregulation of Na_V_ currents is dependent on the trafficking of Na_V_ channels to the surface membrane. This peripheral sensitisation mechanism of DRG neurons can explain the success of macrophage depletion or inhibition strategies to relieve pain *in vivo*. Modulating the electrophysiological state of macrophages by silencing K_ir_2.1 currents is sufficient to prevent the sensitisation effect. Targeting the ion channels and electrophysiological state of non-excitable cells may lead to new avenues in neuropathic pain research.

## Methods

### Animals

Experiments were performed on heterozygote B6.129P-Cx3cr1tm1Litt/J, referred to as CX3CR1-GFP mice (Jackson Laboratory), homozygous TgN(hGFAP-ECFP)-GCED, referred to as hGFAP-CFP mice (a generous gift from Paola Bezzi, Department of Fundamental Neurosciences, University of Lausanne) (52), and wild-type C57BL/6J (Charles-Rivers) adult male and female mice. CX3CR1-GFP transgenic mice have endogenously labelled myeloid cells and are commonly used to label macrophages and microglia (25, 53). Ninety-seven per cent of Iba1^+^ macrophages in DRG express CX3CR1 (10). hGFAP-CFP transgenic mice express CFP in GFAP^+^ glial cells, which include astrocytes, non-myelinating Schwann cells and reactive satellite glial cells (12, 13). The animals were housed in standard cages with free access to food and water at 22 ± 0.5 °C under a controlled 12-hour light/dark cycle. SNI or sham surgery was performed unilaterally under general anaesthesia with isoflurane (Piramal) as described previously (54, 55). Tissue was collected after terminal intraperitoneal injection of pentobarbital (50 mg/kg).

### Western blot

L3–L5 DRG were dissected from mice 7 days after SNI or sham surgery (n = 5 males). The fresh tissue was lysed and protein was separated with sodium dodecyl sulphate–polyacrylamide gel electrophoresis (SDS-PAGE). The separated protein was transferred to polyvinylidene fluoride membranes for western blotting. The primary antibodies were rabbit anti-Iba1 (Wako) and mouse anti-α-tubulin (Sigma-Aldrich), used as a reference protein. The secondary antibodies were horseradish peroxidase–conjugated goat anti-rabbit (Dako) or goat anti-mouse (Dako). The protocol is described in detail in the Supplemental Methods.

### Immunohistochemistry

Age-matched adult mice at 2, 4 and 7 days after SNI, 7 days after sham surgeries and for the naïve condition (n = 4 females and n = 2 males for each time point and condition) were transcardially perfused with saline, followed by 4% paraformaldehyde (PFA, Sigma-Aldrich) in 0.1M phosphate buffer (PB). L3– L5 DRG were collected, post-fixed and cryosectioned at 12 μm for immunostaining using the primary antibodies rabbit anti-Ki67 (Abcam) at 1:500 and rabbit anti-MHC II (Abcam) and the secondary antibody donkey Cy3 anti-rabbit (Jackson). The protocol is described in detail in the Supplemental Methods.

### DRG dissociation

Ipsilateral and contralateral L3–L5 DRG were surgically removed and immersed in cold Hanks’ Balanced Salt Solution (HBSS, Sigma-Aldrich). Subsequently, DRG were incubated for 1 hour at 37 °C with 1 mg/ml dispase II (Roche) and 2 mg/ml collagenase A (Roche) prepared in HBSS. Using a fire polished glass Pasteur pipette, DRG were transferred to 1 ml of warm culture medium: Dulbecco’s Modified Eagle Medium (DMEM, Gibco) supplemented with 10% foetal bovine serum (FBS) (Gibco) and 1% penicillin/streptomycin (Sigma-Aldrich). DRG were then triturated 10–13 times, avoiding bubble formation as much as possible. Four millilitres of culture medium was added to dilute the enzymes and the dissociated DRG were centrifuged at 1000 rpm for 10 min. The supernatant was discarded and the DRG pellet was re-suspended in the appropriate volume depending on the subsequent technical approach.

### Fluorescence-activated cell sorting (FACS) and preparation of conditioned medium (CM)

The dissociated DRG sample was suspended in 1 ml of culture medium and filtered. CX3CR1-GFP and GFAP-CFP cells from adult male and female mice were submitted to FACS immediately after dissociation using a MoFlo Astrios EQ (Beckman Coulter) based on their fluorophore (GFP and CFP). The GFP^+^ and CFP^+^ cells were sorted with a 488 and 405 mm laser, respectively, and collected based on the signal with the 350/30 and 525/50 mm filters, respectively, at room temperature (21–23 °C). Positive events were harvested in 1.5 ml sterile tubes containing 500 μl of culture medium. Subsequently, the cell suspension was diluted in 2.5 ml of culture medium and plated in 35 mm culture dish previously coated for at least 2 hours with 0.1 mg/ml poly-_D_-lysine (Sigma-Aldrich) in the incubator. The culture medium was then collected after 3 days. This was considered to be CX3CR1^+^ cell and GFAP^+^ cell CM throughout the entire study (Figure 1).

To discern the molecular pathways, pharmacological blockers were freshly prepared from stock solutions. CX3CR1^+^ cells were cultured for 24 hours in medium with or without 50 μM brefeldin to block vesicular release. The culture medium was replaced with standard culture medium to prepare and collect CM 3 days later. To inhibit TNF in CM, the DRG culture from naïve mice was pre-treated for 1 hour in normal culture medium with or without 10 μM UCB-9260, and then for 2 hours with 10 μM UCB-9260 diluted in CM. UCB-9260 is a synthetic blocker that directly binds to TNF to prevent its effect on TNF receptors (56). To block p38 in DRG neurons, the DRG culture from naïve mice was pre-treated for 1 hour in normal culture medium with or without 10 μM SB202190, and then for 2 hours with 10 μM SB202190 diluted in CM (26). Nocodazole was used to inhibit transport of proteins to the surface membrane in DRG neurons (50). The DRG culture from naïve mice was pre-treated for 1 hour with medium with or without 10 μM nocodazole, and then for 2 hours with 10 μM SB202190 diluted in normally prepared CM. DRG neurons were patched immediately after the treatments.

### Proteome analysis by LC-MS/MS

CX3CR1^+^ cells from L3-L5 DRG were dissociated 7 days after SNI and sorted by FACS (n = 3 groups containing cells from 5–6 males and 2–3 females, for a total of pooled sample from 8 mice) and centrifuged at 1000 rpm for 10 min to remove the supernatant. The pellets were frozen at −80 °C until proteome analysis by LC-MS/MS. The samples were digested according to a modified version of the iST method, named the miST method (57). Peptides were separated by LC for full MS survey scans. MS/MS data were processed by the MaxQuant software (version 1.6.14) (58) incorporating the Andromeda search engine (59). The details of the protocol and data analysis are found in the Supplemental Methods.

### MEA recordings

DRG from naïve mice were dissociated and cultured overnight on MEA dishes (60MEA200/30iR-Ti-gr, Multi Channel Systems). The dish was coated with 80 μg/ml laminin (Sigma-Aldrich) followed by 0.1 mg/ml poly-_D_-lysine (Sigma-Aldrich). Cells were seeded and placed in the incubator for 1 hour, after which 1 ml of CM was added to each MEA dish. MEA data were collected using a MEA2100-System amplifier (Multi Channel Systems). The extracellular recordings were done for 2 minutes with a frequency of acquisition of 10 000 Hz, immediately after adding CM, after 2 hours of treatment and after overnight treatment. The data collector unit MEA2100-HS head-stage transfers the signal from the MEA dish to the MCS-IFB interface board, which subsequently converts and transfers the digital signal to the main computer. All recordings are controlled by the Multichannel Experimenter software and are done at 37 °C with a TC02 temperature controller. The data analysis was done offline using the Multichannel Analyzer software (Multi Channel Systems), on which a segment of a recording with no activity was filtered using the Butterworth order 2 filter with a high-pass cut-off of 200 Hz and a spike detection threshold of ± 5 standard deviations was used to analyse the entire recording. Afterwards, all files were post-analysed and electrodes with at least five spikes were considered to be active for the final comparison.

### siRNA transfection

siRNA for the mouse gene ID #16518 coding for Kcnj2 (potassium inwardly rectifying channel, subfamily J, member 2) and the siPOOL scramble negative control (siTOOLs Biotech) were used in the experiments (25). In short, 6 μl of INTERFERin reagent for siRNA (Polyplus Transfection) was mixed in 200 μl of serum-free medium containing 50 nM siRNA or scRNA, vortexed and incubated for 10 min at room temperature to allow transfection complexes to form. The mix was then added in the culture medium of FACS CX3CR1^+^ cells and incubated for 24 hours at 5% CO_2_ and 37 °C. The following day, the culture medium was replaced, and CM was collected after 3 days.

### Electrophysiology

#### DRG neurons

After culturing overnight, freshly dissociated DRG from naïve mice were treated with CM for 2 hours. Small DRG neurons (<30 pF in capacitance) were patched in whole-cell conformation with a current clamp protocol consisting of a 500 ms ramp increasing with 50 pA increments from a −60 mV holding potential.

Voltage clamp recording was performed to measure Na_V_ currents. Prior to recording, DRG neurons were held at a –60 mV potential for 5 minutes to dialyse the cell with CsF from the intracellular solution, to reach ion equilibrium and to inactivate the Na_V_1.9 current. The neurons were subsequently clamped at –80 mV for 2 more minutes before starting the recordings.

The steady-state activation (SSA) curves of sodium currents were evoked from a holding potential of −120 mV with 50-ms test pulses ranging from −80 to +70 mV in 5-mV increments. The P/4 procedure was used to digitally subtract the leak currents. The current densities were calculated by dividing the maximum peak amplitude of the current (INa) by the cell’s capacitance. By extrapolating the linear portion of the current–voltage characteristic (I–V) curve through 0 current using the simple linear regression in GraphPad Prism, we determined the reversal potential (Vrev) of each cell. Subsequently, the conductance was calculated as G = I / (Vm – Vrev), where G is conductance, I is the current amplitude, Vm is the current potential and Vrev is the reversal potential. Conductance values from individual cells were fitted with the Boltzmann relationship, G(Vm) = 1 / (1 + exp[(V_½_ – Vm) / k]), where G is the conductance, V_½_ is the voltage at which half of the channels are activated, Vm is the membrane potential and k is the slope factor.

The steady-state inactivation (SSI) curves were measured from a holding potential of −120 mV using a 500-ms pre-pulse to the indicated potentials ranging from −120 to +20 mV in 5-mV increments followed by a 0-mV test pulse. The current densities were normalised to the maximum current densities to subsequently calculate the V_½_ and slope of the SSI for each cell.

Na_V_1.7 and Na_V_1.8 currents were separated as described previously by perfusing the recording chamber with TTX and ProTXII (47). After recording of total Na_V_ currents, 5 nM protoxin II (ProTxII, Sigma-Aldrich) diluted in extracellular solution was perfused for at least 10 min to repeat SSA and SSI recordings. Then, 300 nM tetrodotoxin (TTX, Enzo Life Sciences AG) in extracellular solution was perfused for at least 5 minutes, and SSA and SSI recordings were repeated. The Na_V_1.7 current was analysed by subtracting the current after ProTxII perfusion from the total Na_V_ current. The Na_V_1.8 current was analysed by subtracting the current after TTX perfusion from the total Na_V_ current. For the composition of intracellular and extracellular solutions, see the Supplemental Methods.

#### CX3CR1^+^ cells

Fluorescent cells from the dissociated DRG of CX3CR1-GFP adult male and female mice 7 days after SNI were recorded as described previously (25). They were clamped at –80 mV. Resting membrane potential was measured in 1-minute-long current clamp recording in standard extracellular solution.

Voltage clamp recording was performed in modified extracellular solution with 20 mM KCl to enhance inward-rectifying potassium currents. The composition of intracellular and extracellular solutions can be found in the Supplemental Methods. The voltage clamp protocol consisted of 300-ms-long steps from – 120 mV to +60 mV, increasing in 10-mV increments. Currents were normalised to the capacitance (Cm) before representing the I–V curves. ML133 (Sigma-Aldrich), a K_ir_2.1 channel inhibitor, was diluted to 50 μM in modified extracellular solution from a 50 mM stock solution in dimethyl sulphoxide (DMSO). Recorded GFP^+^ fluorescent cells originated from mice as follows: 10 contralateral cells from 2 males, 19 contralateral cells from 3 females, 14 ipsilateral cells from 2 males, 21 ipsilateral cells from 3 females, 1 ipsilateral cell treated with ML133 from 1 male, and 17 ipsilateral cells treated with ML133 from 2 females.

#### Data acquisition

Data were acquired using a MultiClamp 700B amplifier and a Digidata 1440A driven by the pClamp 10.3 software (Molecular Devices). All whole-cell patch-clamp recordings were made using a BX51WI fixed-stage upright microscope (Olympus) with a CoolLED pE-340fura equipped with a LED eGFP pE-300 filter-set (CoolLED Ltd) and a ORCAFlash2.8 digital camera (Hamamatsu Photonics) under the control of the CellSens v3.2 acquisition software (Olympus). Borosilicate patch pipettes with filament (Sutter Instrument) were pulled using a P-97 pipette puller and fire-polished using a MF200-2 microforge, H4 platinum/iridium wire (World Precision Instruments) and a W30S-LED Revelation III (LW Scientific) at a resistance near 3 MΩ for neurons and 5 MΩ for macrophages. The fast, slow and whole-cell compensations were made with Multiclamp 700B (Molecular Devices) at bandwidth of 5 kHz and a low-pass filter of 10 kHz.

### Statistics

Statistical analyses were done with GraphPad Prism 9.5.1, and the effect size was calculated in R. Two-way analysis of variance (ANOVA) was performed on immunostaining cell counting, MEA, current clamp and voltage clamp data to analyse the relationship between two variables. *Post ho*c multiple comparison analyses were performed for data where the interaction variable was significantly different. When two groups were compared, data were analysed using an unpaired two-tailed Student’s t test. Data are presented as mean ± SEM with individual values or as box-and-whisker plots that show the minimum to maximum values. Proteome data are expressed in a volcano plot indicating the log2 of the fold change and log p value for statistical significance. A p-value <0.05 was considered to be statistically significant. The thresholds were represented as follows: * p <0.05, ** p <0.01, *** p <0.001 and **** p <0.0001. The full list of statistical tests; t, F and p values; and the effect size are provided in Supplemental Table 3. The data presented in this study are freely available from Zenodo.org at doi:10.5281/zenodo.7780850.

### Study approval

All experiments involving animals were approved by the Committee on Animal Experimentation of the Canton de Vaud, Switzerland, in accordance with the Swiss Federal Laws on Animal Welfare and the guidelines of the International Association for the Study of Pain guidelines for the use of animal in research (60).

## Author contribution

EK: designed research, performed research, analysed data, designed the figures, wrote the paper; AD: designed research, performed research, analysed data, wrote the paper; PCSC: supervised the work, designed figures; GK: performed research; ID: designed research, supervised the study, wrote the paper; MS: designed research, supervised the study, wrote the paper.

## Acknowledgements

This study was supported by a grant from the Swiss National Science Foundation (#310030_179169/1 to I. Decosterd, 2018-2022). The authors thank Manfredo Quadroni, Luigi Bozzo, Danny Labes and Mariela Castelblanco for the work performed in the Proteomic Analysis Facility, the Cell Imaging Facility and the Flow Cytometry Facility of University of Lausanne. The schematic drawings were adapted from images available on Servier Medical Art (CC-BY 3.0, smart.servier.com).

## Notes

https://doi.org/10.5281/zenodo.7780850

### Competing Interest Statement

The authors have declared no competing interest.

### Summary of Updates

Figure 4 revised

