## Supplemental Materials for "K_ir_2.1 modulation in macrophages sensitises dorsal root ganglion neurons through TNF secretion after nerve injury"

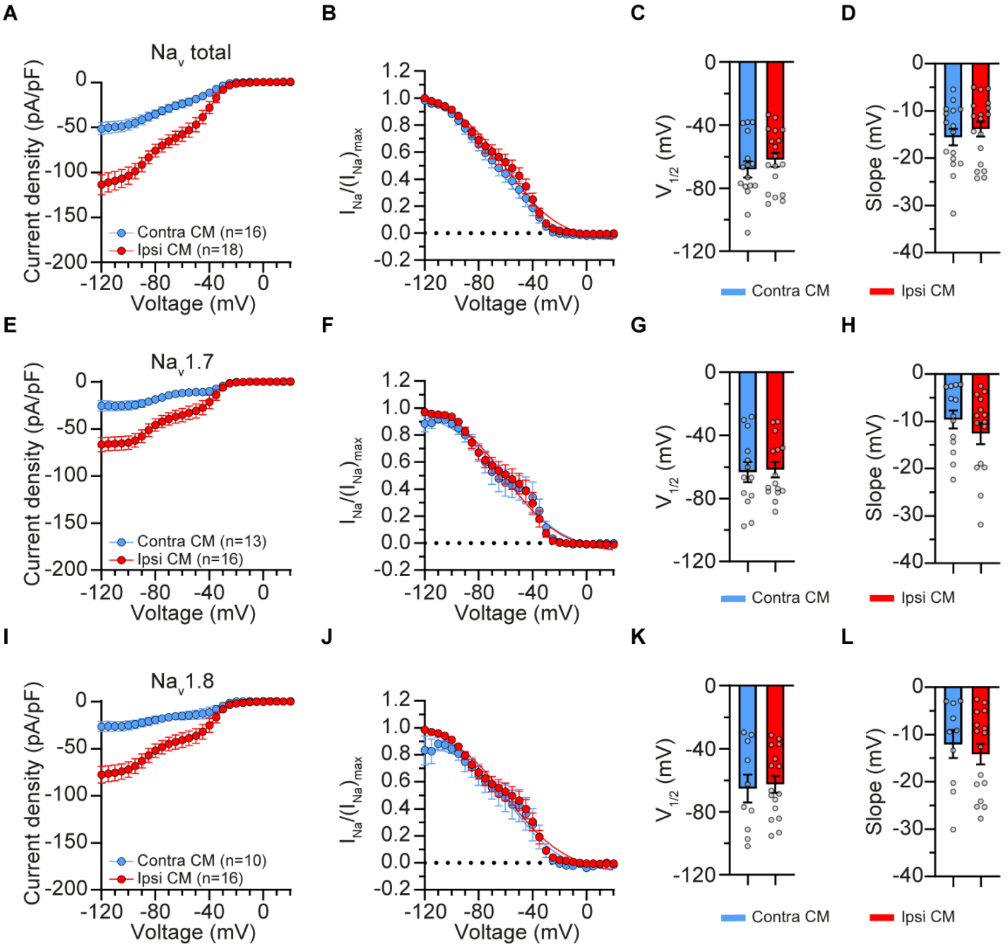

Supplemental figure 1: CX3CR1+ cells conditioned medium (CM) does not affect the steady-state inactivation of total Nav, Nav1.7 and Nav1.8 currents in small dorsal root ganglion (DRG) neurons. Inactivation current densities of total Nav (A), Nav1.7 (E), Nav1.8 (I). Steady state inactivation curves of total Nav (B), Nav1.7 (F), Nav1.8 (J). Half-maximal activation (V½) (C, G, K) and the slope (D, H, L) were calculated for each inactivation curve.

Supplemental table 1: fluorescence-activated cell sorted (FACS) events for conditioned medium (CM) preparation from ipsilateral or contralateral dorsal root ganglia (DRG) from mice after spared nerve injury (SNI).

| FACS events in the CM (events/ml) | Side to SNI | Figures | Experimental groups |
|---|---|---|---|
| 11646<br>7823 | Ipsi<br>Contra | 1BCD | Ipsi vs. contra CX3CR1+ cells CM |
| 13281<br>12195<br>16175<br>14007 | Ipsi<br>Contra<br>Ipsi<br>Contra | 1EFG | Ipsi vs. contra GFAP+ cells CM |
| 3333<br>3333 | Ipsi<br>Contra | 2F<br>5ABCDE<br>FGHIJKLM | Normalized FACS events in CM<br>ipsi vs. contra CX3CR1+ cells CM |
| 5475<br>4600 | Ipsi<br>Ipsi | 4EI | Ipsi CX3CR1+ cells + siRNA Kir2.1<br>vs. ipsi CX3CR1+ cells + scRNA CM |
| 7538<br>3260 | Ipsi<br>Ipsi | 4FGH, JKL<br>5NO | Ipsi CX3CR1+ cells<br>vs. CX3CR1+ cells + drug<br>(Brefeldin, UCB-9260, SB202190 or nocodazole) |

Supplemental table 2: List of significantly regulated proteins measured by liquid chromatography with tandem mass spectrometry (LC-MS/MS) in CX3CR1+ cells from ipsilateral and contralateral dorsal root ganglia (DRG) after spared nerve injury (SNI).

| Gene name | Protein name | Student's T-test Difference ipsi_contra (log2(fold change) | -Log Student's T-test p-value ipsi_contra |
|---|---|---|---|
| Prkab1 | 5-AMP-activated protein kinase subunit beta-1 | 2.986 | 3.95 |
| Smcr8 | Smith-Magenis syndrome chromosomal region candidate gene 8 protein homolog | 2.149 | 3.24 |
| Tgm2 | Protein-glutamine gamma-glutamyltransferase 2 | 1.121 | 3.29 |
| Cd38 | ADP-ribosyl cyclase/cyclic ADP-ribose hydrolase 1 | 1.072 | 3.25 |
| Ppm1f | Protein phosphatase 1F | 0.752 | 3.31 |
| Snx6 | Sorting nexin-6;Sorting nexin-6, N-terminally processed | 0.698 | 3.30 |
| Ctsd | Cathepsin D | 0.536 | 3.40 |
| Cops6 | COP9 signalosome complex subunit 6 | 0.445 | 3.37 |
| Oplah | 5-oxoprolinase | 0.440 | 3.13 |
| Gdi2 | Rab GDP dissociation inhibitor beta | 0.438 | 4.81 |
| Dpysl2 | Dihydropyrimidinase-related protein 2 | 0.412 | 3.78 |
| Eef2 | Elongation factor 2 | 0.381 | 3.26 |
| FAM120A | Constitutive coactivator of PPAR-gamma-like protein 1 | 0.381 | 4.83 |
| Rhot2 | Mitochondrial Rho GTPase 2 | 0.269 | 4.23 |
| Mthfd1 | Methylenetetrahydrofolate dehydrogenase; cyclohydrolase; Formyltetrahydrofolate synthetase 1 | 0.265 | 3.94 |
| Uso1 | General vesicular transport factor p115 | 0.162 | 3.17 |
| Vps33b | Vacuolar protein sorting-associated protein 33B | 0.061 | 3.14 |
| Acad9 | Acyl-CoA dehydrogenase family member 9, mitochondrial | -0.219 | 3.93 |
| Tbl1x | F-box-like/WD repeat-containing protein TBL1X | -0.267 | 3.17 |
| Fam129a | Protein Niban | -0.275 | 3.29 |
| Ddb1 | DNA damage-binding protein 1 | -0.288 | 3.42 |
| Hmg20a | High mobility group protein 20A | -0.362 | 4.39 |
| H2afy | Core histone macro-H2A.1 | -0.470 | 4.09 |
| Gstm1 | Glutathione S-transferase Mu 1 | -0.519 | 5.05 |
| Flna | Filamin-A | -0.556 | 3.15 |
| Myh10 | Myosin-10 | -0.574 | 3.62 |
| Lbr | Lamin-B receptor | -0.590 | 3.98 |
| Engase | Cytosolic endo-beta-N-acetylglucosaminidase | -0.635 | 3.37 |
| Cd74 | H-2 class II histocompatibility antigen gamma chain | -0.694 | 3.66 |
| Ass1 | Argininosuccinate synthase | -0.813 | 3.24 |
| Znf512 | Zinc finger protein 512 | -0.838 | 3.23 |
| Arg1 | Arginase-1 | -2.802 | 3.15 |
| Spen | Msx2-interacting protein | -5.486 | 3.86 |

Supplemental table 3: List of statistical test for significance and effect size for each figure.

| Fig. | Statistical test | N | Stat-value | P-value (Two-tailed) | Effect-size |
|---|---|---|---|---|---|
| **Figure 1** | | | | | |
| 1B | Two-way ANOVA | Contra CM = 154<br>Ipsi CM = 59 | Treatment type $F(1,207) = 12.338$ Treatment duration $F(2,207) = 0.066$ Interaction $F(2,207) = 0.021$ | Treatment type $p < 0.001$ Treatment duration $p = 0.9358$ Interaction $p = 0.9796$ | Treatment type $\eta^2 = 0.056$ Treatment duration $\eta^2 = 0.001$ Interaction $\eta^2 = 0.000$ |
| 1C | Mixed model ANOVA with repeated measures, and post-hoc Sidak comparisons | Contra CM = 10<br>Ipsi CM = 15 | Treatment CM $F(1, 23) = 18.57$ Current $F(2.586, 58.18) = 137.6$ Interaction $F(10, 225) = 6.530$ | Treatment $p = 0.0003$ Current $p < 0.0001$ Interaction $p < 0.0001$ | Treatment $\eta^2 = 0.011$ Current $\eta^2 = 0.816$ Interaction $\eta^2 = 0.039$ |
| 1D | Two-way ANOVA with repeated measures, and post-hoc Sidak comparisons | Contra CM = 15<br>Ipsi CM = 17 | Treatment CM $F(1, 30) = 16.06$ Voltage $F(2.18, 65.38) = 43.73$ Interaction $F(2.18, 65.38) = 9.723$ | Treatment $p = 0.0004$ Voltage $p < 0.0001$ Interaction $p < 0.0001$ | Treatment: $\eta^2_G = 0.187$ Voltage: $\eta^2_G = 0.454$ Interaction: $\eta^2_G = 0.156$ |
| 1E | Two-way ANOVA | Contra CM = 64<br>Ipsi CM = 51 | Treatment $F(1,109) = 0.163$ Treatment duration $F(2,109) = 1.394$ Interaction $F(2,109) = 0.438$ | Treatment type $p = 0.6873$ Treatment duration $p = 0.2525$ Interaction $p = 0.6462$ | Treatment $\eta^2 = 0.000$ Treatment duration $\eta^2 = 0.024$ Interaction $\eta^2 = 0.008$ |
| 1F | Two-way ANOVA with repeated measures | Contra CM = 17<br>Ipsi CM = 11 | Treatment $F(1, 26) = 0.225$ Current $F(2.174, 56.53) = 41.51$ Interaction $F(2.17, 56.53) = 0.848$ | Treatment $p = 0.6393$ Current $p < 0.0001$ Interaction $p = 0.5825$ | Treatment: $\eta^2_G = 0.006$ Current: $\eta^2_G = 0.299$ Interaction: $\eta^2_G = 0.009$ |
| 1G | Two-way ANOVA with repeated measures | Contra CM = 18<br>Ipsi CM = 20 | Treatment $F(1, 36) = 2.851$ Voltage $F(30, 1080) = 68.44$ Interaction $F(30, 1080) = 0.881$ | Treatment $p = 0.1000$ Voltage $p < 0.0001$ Interaction $p = 0.6522$ | Treatment: $\eta^2_G = 0.029$ Voltage: $\eta^2_G = 0.540$ Interaction: $\eta^2_G = 0.015$ |
| **Figure 2** | | | | | |
| 2B | Mixed model ANOVA with repeated measures, and post-hoc Dunnett's comparisons against naïve | Naïve = 6<br>Sham = 6<br>SNI2d = 6<br>SNI4d = 6<br>SNI7d = 6 | Surgery $F(4,25) = 17.96$ DRG $F(1.737, 36.47) = 24.87$ Interaction $(F8,42) = 6.508$ | Surgery $p<0.0001$ DRG $p<0.0001$ Interaction $p<0.0001$ | Surgery $\eta^2 = 0.314$ DRG $\eta^2 = 0.217$ Interaction $\eta^2 = 0.227$ |

| | | | | | |
|---|---|---|---|---|---|
| 2C | Mixed model ANOVA with repeated measures, and post-hoc Dunnett's comparisons against naïve | Naïve = 6<br>Sham = 6<br>SNI2d = 6<br>SNI4d = 6<br>SNI7d = 6 | Surgery F(4,25) = 7.179<br>DRG F(1.791, 34.93) = 16.94 Interaction (F8,39) = 5.289 | Surgery p = 0.0005<br>DRG p<0.0001<br>Interaction p = 0.0002 | Surgery<br>$\eta^2$ = 0.194<br>DRG<br>$\eta^2$ = 0.229<br>Interaction<br>$\eta^2$ = 0.286 |
| 2D | Unpaired student's T-Test, with Welch's correction | Sham = 5,<br>SNI = 5 | t(7.91) = 2.399 | p = 0.0436 | Cohen d = -1.52 |
| 2E | Pearson's Correlation | Trial 1 (9429 cells/ml) = 15,<br>Trial 2 (3337 cells/ml) = 24,<br>Trial 3 (7538 cells/ml) = 11,<br>Trial 4 (7538 cells/ml) = 8,<br>Trial 5 (7538 cells/ml) = 11 | r = 0.935<br><br>$R^2$<br>= 0.874 | p = 0.0199 | |
| 2F | Mixed model ANOVA with repeated measures, and post-hoc Sidak comparisons | Contra CM = 38<br><br>Ipsi CM = 24 | Treatment F(1, 60) = 17.74 Current F(1.851, 102.6) = 101.8<br>Interaction F(10, 554) = 13.33 | Treatment p<0.0001 Current p<0.0001<br>Interaction p<0.0001 | Treatment<br>$\eta^2$ = 0.013<br>Current<br>$\eta^2$ = 0.586<br>Interaction<br>$\eta^2$ = 0.076 |
| **Figure 3** | | | | | |
| 3B | Mixed model ANOVA with repeated measures | Naïve = 4<br>Sham = 4<br>SNI2d = 4<br>SNI4d = 4<br>SNI7d = 4 | Surgery F(4,15) = 2.003<br>DRG F(1.162, 6.973) = 1.367 Interaction F(8, 12) = 1.383 | Surgery p = 0.1458<br>DRG p = 0.2895<br>Interaction p = 0.2945 | Surgery<br>$\eta^2$ = 0.167<br>DRG<br>$\eta^2$ = 0.057<br>Interaction<br>$\eta^2$ = 0.231 |
| 3C | Mixed model ANOVA with repeated measures, and post-hoc Dunnett's comparisons against naïve | Naïve = 4<br>Sham = 4<br>SNI2d = 4<br>SNI4d = 4<br>SNI7d = 4 | Surgery F(4,14) = 18.40<br>DRG F(1.162,6.389) = 21.79 Interaction F(8, 11) = 5.729 | Surgery p < 0.0001<br>DRG p = 0.0025<br>Interaction p = 0.0048 | Surgery<br>$\eta^2$ = 0.400<br>DRG<br>$\eta^2$ = 0.237<br>Interaction<br>$\eta^2$ = 0.249 |
| **Figure 4** | | | | | |
| 4A | Unpaired student's T-Test, with Welch's correction | Contra = 24<br><br>Ipsi = 31 | t(52.57) = 2.996 | p = 0.0042 | Cohen d = -0.78 |

| | | | | | |
|---|---|---|---|---|---|
| 4B | Unpaired student's T-Test, with Welch's correction | Contra = 22<br><br>Ipsi = 31 | t(42.87) = 0.1393 | p = 0.8899 | Cohen d = -0.04 |
| 4C | Two-way ANOVA with repeated measures, and post-hoc Tukey's comparisons | Contra = 12<br><br>Ipsi = 19<br><br>Ipsi + ML133 50µM = 9 | Group (F2,78) = 13.28<br>Voltage (F1.176, 91.54) = 207.1<br>Interaction (48, 1869) = 12.37 | Group p < 0.0001<br>Voltage p<0.0001<br>Interaction p<0.0001 | Group<br>$\eta^2 = 0.004$<br>Surgery<br>$\eta^2 = 0.665$<br>Interaction<br>$\eta^2 = 0.079$ |
| 4E | Mixed model ANOVA with repeated measures, and post-hoc Sidak comparisons | Ipsi CM + Kir2.1 siRNA = 21<br><br>Ipsi CM + scRNA = 15 | Group F(1, 50) = 7.635<br>Current F(2.34,114.4) = 186.8 Interaction F(10,489) = 3.139 | Group p < 0.0001<br>Current p = 0.008<br>Interaction p = 0.0007 | Group<br>$\eta^2 = 0.003$<br>Current<br>$\eta^2 = 0.779$<br>Interaction<br>$\eta^2 = 0.013$ |
| 4F | Mixed model ANOVA with repeated measures, and post-hoc Sidak comparisons | Ipsi CM = 14<br><br>Ipsi CM + Brefeldin = 16 | Treatment F(1,28) = 8.270 Current F(1.406,37.26) = 69.99 Interaction F(10,265) = 9.285 | Treatment p = 0.0063 Current p < 0.0001 Interaction p < 0.0001 | Treatment<br>$\eta^2 = 0.008$<br>Current<br>$\eta^2 = 0.656$<br>Interaction<br>$\eta^2 = 0.087$ |
| 4G | Mixed model ANOVA with repeated measures, and post-hoc Sidak comparisons | Ipsi CM = 12<br><br>Ipsi CM + UCB = 15 | Treatment F(1,25) = 14.88 Current F(2.260,49.95) = 53.53 Interaction F(10,221) = 10.80 | Treatment p = 0.0007 Current p < 0.0001 Interaction p < 0.0001 | Treatment<br>$\eta^2 = 0.017$<br>Current<br>$\eta^2 = 0.608$<br>Interaction<br>$\eta^2 = 0.123$ |
| 4H | Mixed model ANOVA with repeated measures, and post-hoc Sidak comparisons | Ipsi CM = 8<br><br>Ipsi CM + SB = 11 | Treatment F(1,17) = 22.10 Current F(10, 163) = 43.46 Interaction F(10,163) = 18.71 | Treatment p = 0.0002 Current p < 0.0001 Interaction p < 0.0001 | Treatment<br>$\eta^2 = 0.027$<br>Current<br>$\eta^2 = 0.538$<br>Interaction<br>$\eta^2 = 0.232$ |
| 4I | Two-way ANOVA with repeated measures, and post-hoc Sidak comparisons | Ipsi CM + Kir2.1 siRNA = 18<br><br>Ipsi CM + scRNA = 10 | Treatement F(1,26) = 10.09 Voltage F(1.195, 31.06) = 29.74 Interaction F(30, 780) = 8.075 | Treatment p = 0.0038 Voltage p < 0.0001 Interaction p < 0.0001 | Treatment: $\eta^2_G$= 0.147<br>Voltage: $\eta^2_G$= 0.389<br>Interaction: $\eta^2_G$= 0.148 |
| 4J | Mixed model ANOVA with repeated measures, and post-hoc Sidak comparisons | Ipsi CM = 9<br><br>Ipsi CM + Brefeldin = 8 | Treatment F(1,15) = 8.303 Voltage F(1.229,18.22) = 25.39 Interaction F(30,445) = 4.1 | Treatment p = 0.0114 Voltage p < 0.0001 Interaction p < 0.0001 | Treatment<br>$\eta^2 = 0.006$<br>Voltage<br>$\eta^2 = 0.569$<br>Interaction<br>$\eta^2 = 0.092$ |

| | | | | | |
|---|---|---|---|---|---|
| 4K | Two-way ANOVA with repeated measures, and post-hoc Sidak comparisons | Ipsi CM = 13<br><br>Ipsi CM + UCB = 12 | Treatment $F(1,23) = 9.021$ Voltage $F(1.415,32.55) = 20.40$ Interaction $F(30,690) = 6.835$ | Treatment $p = 0.0063$ Voltage $p < 0.0001$ Interaction $p < 0.0001$ | Treatment: $\eta^2_G$= 0.140<br>Voltage: $\eta^2_G$= 0.341<br>Interaction: $\eta^2_G$= 0.148 |
| 4L | Two-way ANOVA with repeated measures, and post-hoc Sidak comparisons | Ipsi CM = 7<br><br>Ipsi CM + SB = 12 | Treatment $F(1,17) = 46.59$ Voltage $F(2.310, 39.27) = 69.09$ Interaction $F(30,510) = 19.44$ | Treatment $p < 0.0001$ Voltage $p < 0.0001$ Interaction $p < 0.0001$ | Treatment: $\eta^2_G$= 0.446<br>Voltage: $\eta^2_G$= 0.742<br>Interaction: $\eta^2_G$= 0.447 |
| **Figure 5** | | | | | |
| 5A | Two-way ANOVA with repeated measures, and post-hoc Sidak comparisons | Contra CM = 18<br><br>Ipsi CM = 19 | Treatment $F(1,35) = 30.44$ Voltage $F(2.793,97.77) = 40.42$ Interaction $F(30,1050) = 6.379$ | Treatment $p < 0.0001$ Voltage $p < 0.0001$ Interaction $p < 0.0001$ | Treatment: $\eta^2_G$= 0.180<br>Voltage: $\eta^2_G$= 0.463<br>Interaction: $\eta^2_G$= 0.120 |
| 5C | Unpaired student's T-Test, with Welch's correction | Contra CM = 14<br><br>Ipsi CM = 11 | $t(12.89) = 4.579$ | $p < 0.0001$ | Cohen d = 2.01 |
| 5D | Unpaired student's T-Test, with Welch's correction | Contra CM = 14<br><br>Ipsi CM = 11 | $t(22.94) = 2.063$ | $p = 0.0507$ | Cohen d = 0.80 |
| 5E | Two-way ANOVA with repeated measures, and post-hoc Sidak comparisons | Contra CM = 17<br><br>Ipsi CM = 19 | Treatment $F(1,34) = 26.31$ Voltage $F(2.833,96.31) = 14.34$ Interaction $F(30,1020) = 4.490$ | Treatment $p < 0.0001$ Voltage $p < 0.0001$ Interaction $p < 0.0001$ | Treatment: $\eta^2_G$= 0.146<br>Voltage: $\eta^2_G$= 0.247<br>Interaction: $\eta^2_G$= 0.093 |
| 5G | Unpaired student's T-Test, with Welch's correction | Contra CM =12<br><br>Ipsi CM = 8 | $t(8.11) = 2.501$ | $p = 0.0365$ | Cohen d = 1.35 |
| 5H | Unpaired student's T-Test, with Welch's correction | Contra CM =12<br><br>Ipsi CM = 8 | $t(18) = 1.808$ | $p = 0.0874$ | Cohen d = 0.76 |

| | | | | | |
|---|---|---|---|---|---|
| 5I | Two-way ANOVA with repeated measures, and post-hoc Sidak comparisons | Contra CM = 13<br><br>Ipsi CM = 16 | Treatment $F(1,27) = 28.42$ Voltage $F(2.594, 70.04) = 17.13$ Interaction $F(30, 810) = 4.193$ | Treatment $p < 0.0001$ Voltage $p < 0.0001$ Interaction $p < 0.0001$ | Treatment: $\eta^2_G = 0.183$<br>Voltage: $\eta^2_G = 0.333$<br>Interaction: $\eta^2_G = 0.109$ |
| 5K | Unpaired student's T-Test, with Welch's correction | Contra CM =12<br><br>Ipsi CM = 11 | $t(20.64) = 1.1308$ | $p = 0.2711$ | Cohen d = 0.47 |
| 5L | Unpaired student's T-Test, with Welch's correction | Contra CM =11<br><br>Ipsi CM = 11 | $t(19.99) = 1.3775$ | $p = 0.1836$ | Cohen d = 0.59 |
| 5N | Mixed model ANOVA with repeated measures, and post-hoc Sidak comparisons | Ipsi CM = 9<br><br>Ipsi CM + Nocodazole = 15 | Treatment $F(1,22) = 14.92$ Current $F(1.742, 37.29) = 67.39$ Interaction $F(10, 214) = 8.317$ | Treatment $p = 0.0008$ Current $p < 0.0001$ Interaction $p < 0.0001$ | Treatment $\eta^2 = 0.015$<br>Current $\eta^2 = 0.683$<br>Interaction $\eta^2 = 0.084$ |
| 5O | Mixed model ANOVA with repeated measures, and post-hoc Sidak comparisons | Ipsi CM = 13<br><br>Ipsi CM + Nocodazole = 10 | Treatment $F(1,21) = 16.13$ Current $F(2.125, 44.63) = 23.87$ Interaction $F(30, 630) = 6.882$ | Treatment $p = 0.0006$ Current $p < 0.0001$ Interaction $p < 0.0001$ | Treatment: $\eta^2_G = 0.220$<br>Voltage: $\eta^2_G = 0.419$<br>Interaction: $\eta^2_G = 0.172$ |
| **Supplemental Figure 1** | | | | | |
| 1A | Two-way ANOVA with repeated measures, and post-hoc Sidak comparisons | Contra CM =16<br><br>Ipsi CM = 18 | Treatment $F(1,32) = 23.61$ Voltage $F(2.010, 64.33) = 106.1$ Interaction $F(28, 896) = 14.88$ | Treatment $p < 0.0001$ Voltage $p < 0.0001$ Interaction $p < 0.0001$ | Treatment: $\eta^2_G = 0.246$<br>Voltage: $\eta^2_G = 0.649$<br>Interaction: $\eta^2_G = 0.206$ |
| 1C | Unpaired student's T-Test, with Welch's correction | Contra CM =16<br><br>Ipsi CM = 18 | $t(30.72) = 0.9168$ | $p = 0.3664$ | Cohen d = -0.32 |
| 1D | Unpaired student's T-Test, with Welch's correction | Contra CM =16<br><br>Ipsi CM = 18 | $t(31.201) = 0.7157$ | $p = 0.4795$ | Cohen d = -0.25 |

| | | | | | |
|---|---|---|---|---|---|
| 1E | Two-way ANOVA with repeated measures, and post-hoc Sidak comparisons | Contra CM =13<br><br>Ipsi CM = 16 | Treatment F(1,27) = 13.34 Voltage F(2.262,61.08) = 53.53 Interaction F(28, 756) = 10.61 | Treatment p = 0.0011 Voltage p < 0.0001 Interaction p < 0.0001 | Treatment: $\eta^2_G$= 0.196 Voltage: $\eta^2_G$= 0.501 Interaction: $\eta^2_G$= 0.166 |
| 1G | Unpaired student's T-Test, with Welch's correction | Contra CM =13<br><br>Ipsi CM = 15 | t(23.243) = 0.1684 | p = 0.8677 | Cohen d = -0.06 |
| 1H | Unpaired student's T-Test, with Welch's correction | Contra CM =13<br><br>Ipsi CM = 15 | t(25.861) = 1.002 | p = 0.3254 | Cohen d = 0.37 |
| 1I | Two-way ANOVA with repeated measures, and post-hoc Sidak comparisons | Contra CM =10<br><br>Ipsi CM = 16 | Treatment F(1,24) = 11.89 Voltage F(2.120, 50.87) = 44.73 Interaction F(28, 672) = 10.21 | Treatment p = 0.0021 Voltage p < 0.0001 Interaction p < 0.0001 | Treatment: $\eta^2_G$= 0.206 Voltage: $\eta^2_G$= 0.470 Interaction: $\eta^2_G$= 0.168 |
| 1K | Unpaired student's T-Test, with Welch's correction | Contra CM =10<br><br>Ipsi CM = 16 | t(15.713) = 0.2559 | p = 0.8014 | Cohen d = -0.11 |
| 1L | Unpaired student's T-Test, with Welch's correction | Contra CM =10<br><br>Ipsi CM = 16 | t(18.416) = 0.569 | p = 0.5762 | Cohen d = 0.23 |

Supplemental table 4: List of reagents, solutions, and RRID references used in the current study.

| Reagent or resource | Source (Company, City, State, Country) | Catalogue Number | RRID (Resource Identification Portal) |
|---|---|---|---|
| **Mice** | | | |
| Mouse line CX3CR1-eGFP | Jackson Laboratory, Bar Harbor, ME, USA | B6.129P-Cx3cr1tm1Litt/J, #005582 | IMSR_JAX:005582 |
| Mouse line GFAP-CFP | Frank Kirchhoff, Max Planck Institute of Experimental Medicine, Göttingen, Germany | TgN(hGFAP-ECFP)-GCED | |
| Mouse line C57Bl6j | Charles-Rivers, Wilmington, MA, USA | strain code 680, C57Bl6j | |
| **Materials / Equipment** | | | |
| PVDF membrane | BioRad, Hercules, California (CA), USA | #162-0177 | |
| LAS4000 Fuji system (ImageQuant LAS 4000) | GE Healthcare, Chicago, Illinois (IL), USA | LAS4000 | |
| Zeiss Axio Scan.Z1 slide scanner | Zeiss, Oberkochen, Germany | | |
| Falcon 5 mL Round Bottom Polystyrene Test Tube, with Cell Strainer Snap Cap | Corning, Corning, NY, USA | 352235 | |
| MoFlo Astrios EQ | Beckman Coulter, Brea, CA, USA | #B25982 | |
| Strong cation exchange (SCX) plate (Oasis MCX) | Waters Corp., Milford, MA | | |
| Exploris 480 Orbitrap mass spectrometer | Thermo Fisher Scientific Inc., Waltham, Massachusetts (MA), USA | | |
| Ultimate 3000 RSLCnano HPLC system | Dionex, Sunnyvale, California, US | | |
| Reversed-phase custom packed 40 cm C18 column (75 µm ID, 100Å, Reprosil Pur 1.9 um particles) | Dr. Maisch, Ammerbuch, Germany | | |
| MEA dishes (60MEA200/30iR-Ti-gr) | Multi Channel Systems, Reutlingen, Germany | #890103 | |
| ImmEdge Hydrophobic Barrier PAP Pen | Vector Laboratories, Burlingame, CA, USA | #H-4000 | |
| MEA2100-System amplifier | Multi Channel Systems, Reutlingen, Germany | MEA2100-System amplifier | |
| Thermanox plastic coverslips | Nunc, NY, Rochester | #174950 | |
| Multiclamp Axon Amplifier 700B | Molecular Devices, San Jose, California (CA), USA | #700B | |
| Digidata 1440A | Molecular Devices, San Jose, California (CA), USA | #1440A | |
| CoolLED pE-340 fura | CoolLED Ltd, Andover, UK | #pE-340 fura | |
| LED eGFP pE-300 filterset | CoolLED Ltd, Andover, UK | #E3990113, Exciter: 460/30; Emitter: 520/40 | |
| ORCAFlash2.8, model C11440-10C | Hamamatsu Photonics, Shizuoka, Japan | #820504 | |

| | | | |
|---|---|---|---|
| Olympus BX63 fluorescent microscope | Olympus, Tokyo, Japan | BX63 | |
| TH4 halogen lamp power supply unit | Olympus, Tokyo, Japan | TH4 | |
| MF200-2 microforge | World Precision Instruments, Sarasota, FL, USA | MF200-2 | |
| H4 platinum/iridium wire | World Precision Instruments, Sarasota, FL, USA | MF-200 H4 | |
| Borosilicate patch pipettes with filament | Novato, CA, USA | #BF150-86-7.5HP | |
| W30S-LED Revelation III | LW Scientific, Lawrenceville, GA, USA | W30S | |
| Perfusion valve controller VC-6 | Warner Instrument, Hamden, CT, USA | VC-6 | |
| Gilson minipuls suction system | Gilson, Middleton, WI, USA | | |
| Nalgene 4 mm syringe filters | Thermo Fisher Scientific Inc., Waltham, Massachusetts (MA), USA | 176-0020 | |
| SevenCompact S210 | Mettler Toledo, Columbus, OH, USA | S210 | |
| Osmometer 3320 | Advanced Instruments Inc | 3320 | |
| **Software** | | | |
| Fiji (ImageJ Software) | Schindelin et al., 2012 | | SCR_002285 |
| Zeiss Blue software | Zeiss, Oberkochen, Germany | | SCR_013672 |
| Xcalibur 4.2 software | Thermo Fisher Scientific Inc., Waltham, Massachusetts (MA), USA | | |
| MaxQuant software (version 1.6.14) | Cox and Mann, 2008 | | |
| Andromeda search engine | Cox et al, 2011 | | |
| Perseus software | Tyanova et al, 2016 | | SCR_015753 |
| Multichannel Analyzer | Multi Channel Systems, Reutlingen, Germany | | |
| CellSens v3.2 | Olympus, Tokyo, Japan | | |
| pClamp 10 Software | Molecular Devices, San Jose, California (CA), USA | | SCR_011323 |
| **Plasmids** | | | |
| Silencing RNA for Kir2.1 | Polyplus transfection | | |
| siPOOL scramble RNA | siTOOLs Biotech | | |
| **Antibodies** | | | |
| Rabbit anti-iba1 | Wako Pure Chemical Industries Ltd, Richmond, VA, USA | 19741 | AB_839504 |
| Rabbit anti-α-tubulin | Sigma-Aldrich Inc., Saint-Louis, Missouri (MO), USA | #T5168 | AB_477579 |
| HRP-linked goat anti-rabbit | Aligent, Santa Clara, CA | P0448 | AB_2617138 |
| HRP-linked goat anti-mouse | Aligent, Santa Clara, CA | P0447 | AB_2617137 |
| Rabbit anti-Ki67 | Abcam, Waltham, MA, USA | ab9260 | |
| Rabbit anti-MHC class II | Abcam, Waltham, MA, USA | ab245692 | |
| Donkey Cy3 anti-rabbit | Jackson ImmunoResearch, Ely, UK | 711-165-152 | AB_2307443 |
| **Reagents** | | | |
| Isoflurane | Piramal, Mumbai, Maharashtra, India | G45C19A | |
| Pentobarbital | Streuli Pharma, USA | V102013 | |

| | | |
|---|---|---|
| Hanks' Balanced Salt solution | Sigma-Aldrich Inc., Saint-Louis, Missouri (MO), USA | H6648-500ML |
| Tris | AppliChem, Darmstadt, Germany | #A1086 |
| NaCl 150 mM | Polyplus transfection, New-York, New-York (NY), USA | #702-50 |
| Complete protease inhibitor cocktail | Roche, Basel, Switzerland | #11697498001 |
| PhenylMethaneSulfonyl Fluoride (PMSF) | Sigma-Aldrich Inc., Saint-Louis, Missouri (MO), USA | #P7626 |
| Triton X-100 | Sigma-Aldrich Inc., Saint-Louis, Missouri (MO), USA | #T9284 |
| SDS (Sodium Dodecyl Sulfate) | Sigma-Aldrich Inc., Saint-Louis, Missouri (MO), USA | #L4390 |
| Bovine Serum Albumin | Sigma-Aldrich Inc., Saint-Louis, Missouri (MO), USA | A3294 |
| Protein Assay Dye Reagent Concentrate | Bio-Rad, Hercules, CA, USA | 500-0006 |
| SuperSignal West Dura Extended Duration Substrate | Thermo Fisher Scientific Inc., Waltham, Massachusetts (MA), USA | #34075 |
| Paraformaldehyde | Sigma-Aldrich Inc., Saint-Louis, Missouri (MO), USA | P6148 |
| Phosphate Buffered Saline | Bischel AG, Bern, Switzerland | 100 0 325 |
| Sucrose (D+ Saccharose) | AppliChem, Darmstadt, Germany | #A2211 |
| Tissue-Tek O.C.T. Compound | Sakura Finetek, Alphen aan den Rijn, Netherlands | #4583 |
| Normal goat serum | Vector Laboratories, Newark, CA, USA | S-1000 |
| Normal horse serum | Vector Laboratories, Newark, CA, USA | S-2000 |
| Mowiol 4–88 medium | Calbiochem, San Diego, CA, USA | #475904 |
| Dispase II | Roche, Basel, Switzerland | 4942078001 |
| Collagenase A | Roche, Basel, Switzerland | 10103578001 |
| DMEM | Gibco, Billings, MT, USA | 41965-039 |
| Fetal Bovine Serum | Gibco, Billings, MT, USA | 10082 |
| Penicillin/Streptomycin | Sigma-Aldrich Inc., Saint-Louis, Missouri (MO), USA | P0781 |
| Poly-D-Lysine | Sigma-Aldrich Inc., Saint-Louis, Missouri (MO), USA | P0899 |
| Benzonase | Merck Millipore, Burlington, Massachusetts (MA), USA | #70746 |
| Trypsin/LysC mix | Promega, Madison, WI, USA | #V5073 |
| Laminin | Sigma-Aldrich Inc., Saint-Louis, Missouri (MO), USA | #L2020 |
| INTERFERin reagent for siRNA | Polyplus transfection, France | #89129 |
| Protoxin II (ProTxII) | Sigma-Aldrich Inc., Saint-Louis, Missouri (MO), USA | #P0033 |
| Tetrodotoxin (TTX) | Enzo Life Sciences AG, Farmingdale, NY, USA | 630-002-M001 |
| Brefeldin | Sigma-Aldrich Inc., Saint-Louis, Missouri (MO), USA | B6542-5MG |
| UCB-9260 | MedChemExpress, Monmouth Junction, NJ, USA | HY-133122 |
| SB202190 | Sigma-Aldrich Inc., Saint-Louis, Missouri (MO), USA | S7067-5MG |

| | | | |
|---|---|---|---|
| Nocodazole | Sigma-Aldrich Inc., Saint-Louis, Missouri (MO), USA | M1404-10MG | |
| DMSO | Sigma-Aldrich Inc., Saint-Louis, Missouri (MO), USA | D-2650 | |
| ML133 | Sigma-Aldrich Inc., Saint-Louis, Missouri (MO), USA | SML0190-5MG | |
| NaCl (Sodium Chloride) | Sigma-Aldrich Inc., Saint-Louis, Missouri (MO), USA | #S9625 | |
| CaCl2 (Calcium chloride dihydrate) | Merck Millipore, Burlington, Massachusetts (MA), USA | #102382 | |
| MgCl2-6H2O (Magnesium chloride hexahydrate) | Merck Millipore, Burlington, Massachusetts (MA), USA | #105832 | |
| HEPES | AppliChem, Darmstadt, Germany | #A1069 | |
| Glucose D-(+) | Sigma-Aldrich Inc., Saint-Louis, Missouri (MO), USA | #G7021 | |
| Cd-Cl2 (Cadmium chloride hydrate) | Sigma-Aldrich Inc., Saint-Louis, Missouri (MO), USA | #529575 | |
| TEA-Cl (Tetraethylammonium chloride) | Sigma-Aldrich Inc., Saint-Louis, Missouri (MO), USA | #T2265 | |
| CsF (Cesium fluoride) | Sigma-Aldrich Inc., Saint-Louis, Missouri (MO), USA | #289345 | |
| EGTA (Ethylene Glycol Tetra Acetic Acid) | Sigma-Aldrich Inc., Saint-Louis, Missouri (MO), USA | #E4378 | |
| Cs-OH (Cesium hydroxide monohydrate) | Sigma-Aldrich Inc., Saint-Louis, Missouri (MO), USA | #516988 | |
| KCl (Potassium Chloride) | Sigma-Aldrich Inc., Saint-Louis, Missouri (MO), USA | #P9333 | |
| NaOH (Sodium hydroxide) | Sigma-Aldrich Inc., Saint-Louis, Missouri (MO), USA | #S8045 | |
| Mg-ATP | Sigma-Aldrich Inc., Saint-Louis, Missouri (MO), USA | #A-9187 | |
| KOH | Sigma-Aldrich Inc., Saint-Louis, Missouri (MO), USA | P5958-250G | |
| MgCl2-6H2O (Magnesium chloride hexahydrate) | Merck Millipore, Burlington, Massachusetts (MA), USA | #105832 | |
| Choline-Cl | Sigma-Aldrich Inc., Saint-Louis, Missouri (MO), USA | #C1879 | CAS 67-48-1 |

### Supplemental Methods

#### Western blot

L3-L5 DRG ipsilateral and contralateral to the surgery were dissected out in cold HBSS solution (Sigma-Aldrich) after mice were terminally injected with pentobarbital (50 mg/kg). The fresh DRG tissue were lysed in 100 mM Tris HCl pH 6.8, 2% SDS, 10% glycerol, Complete Protease inhibitor cocktail tablets (Roche). Soluble fractions were recovered in supernatants after 20 min of centrifugation at 10 000 rpm at 4 °C. Protein concentration was measured using Bradford BSA standard scale (Sigma-Aldrich) with Assay dye (Bio-rad). Proteins were separated on acrylamide SDS-PAGE and then transferred to polyvinylidene fluoride membranes (Bio-Rad) that were immunoblotted with the following antibodies: rabbit anti-iba1 (1:1000 in milk; 19741, Wako), and mouse anti-α-tubulin used as a reference protein (1:20,000 in milk; Sigma-Aldrich). We used secondary HRP-linked goat anti-rabbit (1:10 000 in milk, P0448, Dako) or goat anti-mouse (1:10 000 in milk, P0447, Dako) and SuperSignal West Dura Extended Duration Substrate (Thermo Fisher Scientific) for detection. Chemiluminescence was detected by using an imaging system (model LAS-4000 Imaging System; ImageQuant, GE FujiFilm) coupled with an integrated CCD camera. Protein quantification was performed using ImageJ software (Fiji; (1). The quantified signals for the proteins of interest were normalized to reference protein signals.

#### Immunohistochemistry

Mice were terminally anesthetized by i.p injection of pentobarbital and transcardially perfused with saline, followed by 4% paraformaldehyde (PFA, Sigma-Aldrich) in 0.1M phosphate buffer (PB). L3-L5 DRGs were dissected and post-fixed at 4°C for 90 min and then transferred in 20% sucrose in 0.1M PB for 24h. The DRG were rapidly frozen in embedding solution (Tissue-Tek® O.C.T. Compound, Sakura® Finetek) to be cut in 12um thickness section with a cryostat, directly onto slides for immunostaining. After washing in PBS, the sections were incubated in blocking solution of 10% normal goat serum (NGS) (Vector) or normal horse serum (NHS) (Vector), 0.3% triton X-100 (Sigma-Aldrich) in PBS for 30min at room temperature (RT). The following primary antibodies were diluted in 5% NGS or NHS in 0.1% triton X-100 and incubated overnight at 4°C: rabbit anti-Ki67 at 1:500 (Abcam), rabbit anti-MHC class II at 1:1000 (Abcam). After 3 washed in PBS, the sections were incubated in donkey Cy3 anti-rabbit at 1:400 (Jackson ImmunoResearch) secondary antibodies diluted in 1% NGS or 1% NHS in 0.1% triton X-100 for 2 hours at RT. After three final washes in PBS, the sections were dried and covers were mounted with Mowiol 4–88 medium (Calbiochem). Epifluorescence images of the DRG sections were acquired at x20 magnification with Zeiss Axio Scan.Z1 slide scanner using the Zeiss Blue software. Cell counting were performed using the Cell Counter Plugin in ImageJ software (Fiji).

**Proteome analysis by liquid chromatography-tandem mass spectrometry (LC-MS/MS)**

Replicate samples were digested according to a modified version of the iST method (2) (named miST method). Briefly, frozen cell pellets were resuspended in 30ul miST lysis buffer (1% sodium deoxycholate, 100mM Tris pH 8.6, 10 mM DTT ) by vortexing. Resuspended samples were heated at 95°C for 5min. Samples were then diluted 1:1 (v:v) with water containing 4mM $MgCl_2$ and Benzonase (Merck #70746, 100x dil of stock = 250 Units/ul) and incubated for 15 minutes at RT to digest nucleic acids. Reduced disulfides were alkylated by adding ¼ vol (25 ul) of 160 mM chloroacetamide (final 32 mM) and incubating at 25°C for 45min in the dark. Samples were adjusted to 3 mM EDTA and digested with 0.2 ug Trypsin/LysC mix (Promega #V5073) for 1h at 37°C, followed by a second 1h digestion with a second and identical aliquot of proteases. To remove sodium deoxycholate, two sample volumes of isopropanol containing 1% TFA were added to the digests, and the samples were desalted on a strong cation exchange (SCX) plate (Oasis MCX ; Waters Corp., Milford, MA) by centrifugation. After washing with isopropanol/1%TFA, peptides were eluted in 250ul of 80% MeCN, 19% water, 1% (v/v) ammonia.

Eluates after SCX desalting were dried, and resuspended in 20 ul of 0.05% trifluoroacetic acid, 2% acetonitrile. Variable volumes (4-8ul) were injected to equilibrate loaded amounts, based on numbers of sorted cells. Data-dependent LC-MS/MS analysis of was carried out on an Exploris 480 Orbitrap mass spectrometer (Thermo Fisher Scientific) interfaced through a nano-electrospray ion source to an Ultimate 3000 RSLCnano HPLC system (Dionex). Peptides were separated on a reversed-phase custom packed 40 cm C18 column (75 μm ID, 100Å, Reprosil Pur 1.9 um particles, Dr. Maisch, Germany) with a 4-76% acetonitrile gradient in 0.1% formic acid (total time 140 min). Full MS survey scans were performed at 120'000 resolution. A data-dependent acquisition method controlled by Xcalibur 4.2 software (Thermo Fisher Scientific) was used that optimized the number of precursors selected ("top speed") of charge $2^+$ to $5^+$ in the m/z window 375-1500, while maintaining a fixed scan cycle of 2.0s. The precursor isolation window used was 1.6 Th. HCD fragmentation mode was used at a normalized collision energy of 30% and MS/MS spectra were acquired at 15’000 resolution. Peptides selected for MS/MS were excluded from further fragmentation during 60s.

**Mass spectrometry data analysis**

Tandem MS data were processed by the MaxQuant software (version 1.6.14) (3) incorporating the Andromeda search engine (4). The UniProt *Mus musculus* reference proteome (RefProt) database of August 26$^{th}$, 2020 was used (55’508 sequences), supplemented with sequences of common contaminants. Trypsin (cleavage at K,R) was used as the enzyme definition, allowing 2 missed cleavages. Carbamidomethylation of cysteine was specified as a fixed modification. N-terminal acetylation of protein and oxidation of methionine were specified as variable modifications. All identifications were filtered at 1% FDR at both the peptide and protein levels with default MaxQuant parameters. MaxQuant data were further processed with Perseus software (5). iBAQ (6) values were used for quality control assessment, after which LFQ values (7) were used for quantitation after log2 transformation.

Only protein groups were retained which had all three valid values in at least one condition. In the resulting table, missing values were imputed with values determined from a normal distribution (MaxQuant default parameters, width 0.3 and down-shift by 2.2 SD).  A student’s T-test was performed to find significantly different proteins between the two conditions. Multiple testing correction by the permutation method was used to adjust p-values and a threshold of 0.05 was set on the corrected p-values (q-values).

GO and KEGG annotation terms were added. 1D enrichment on the fold-change between the two conditions was performed as described (8), with FDR correction using the Benjamini-Hochberg method and threshold at q=0.02. The final table was filtered to retain only proteins quantified globally by a minimum of two razor or unique peptides.

**Solutions for whole-cell patch clamp**

For sodium recording in DRG neurons an extracellular solution (ES) was used containing (in mM) 30 NaCl, 3 KCl, 1 CaCl2*2H2O, 1 MgCl2*6H2O, 10 HEPES, 10 glucose, 0.1 CdCl, 110 TEA-Cl, and the pH was adjusted to 7.4 with Tris base and the osmolarity was set between 305-310 mOsm with glucose.

The internal solution for sodium recordings contained (in mM) 10 NaCl, 140 CsF, 0.1 CaCl2*2H2O, 0.1 MgCl2*6H2O, 10 HEPES, 1.1 EGTA in CsOH, and the pH was adjusted to 7.35 with CsOH and the osmolarity was set between 290-300 mOsm with glucose. On the other side, for current clamp recordings of APs in DRG neurons the ES solution contained (in mM) 140 NaCl, 3 KCl, 2 CaCl2*2H2O, 2 MgCl2*6H2O, 10 HEPES, and the pH was adjusted to 7.4 with NaOH and the osmolarity was set between 305-310 mOsm with glucose. The IS solution had (in mM) 140 KCl, 3 Mg-ATP, 0.5 EGTA, 5 HEPES, 10 glucose, and the pH was adjusted to 7.35 with KOH and the osmolarity was set between 290-300 mOsm with glucose. The ES to record resting membrane potential of DRG macrophages contained (in mM) 120 NaCl, 5 KCl, 2 CaCl2, 1 MgCl2, 10 HEPES, 10 D-glucose, 15 ChoCl and the pH was adjusted to 7.4 with NaOH and the osmolarity was set between 305-310 mOsm with glucose. Modified ES solution was used to record Kir2.1 currents, and contained (in mM) 120 NaCl, 20 KCl, 2 CaCl2, 1 MgCl2, 10 HEPES, 10 D-glucose, and the pH was adjusted to 7.4 with NaOH and the osmolarity was set between 305-310 mOsm with glucose. The IS contained (in mM) 5 NaCl, 130 KCl, 1 CaCl2, 2 MgCl2, 10 HEPES, 10 EGTA, and the pH was adjusted to 7.35 with KOH and the osmolarity was set between 290-300 mOsm with glucose. IS were filtered using Nalgene 4 mm syringe filters (176-0020, Thermo). A SevenCompact™ S210 (Mettler Toledo) was used for adjusting the pH of solutions and an Osmometer 3320 (Advanced Instruments Inc) was used to calibrate the osmolarity.
